# Real-world longitudinal multi-omics separates weight-loss-associated responses to incretin agonists from a broader response not dependent on weight loss

**DOI:** 10.64898/2026.11.13.766211

**Authors:** T Long, A Large, K Gallagher, JL Justice, K Dao, S Park, J Usuka, J Watrous, M Jain

**Affiliations:** Sapient Bioanalytics, LLC; 10421 Wateridge Circle, Suite 100; San Diego, CA 92121

**Author notes:** Address for correspondence: Mohit Jain, MD PhD Tao Long, PhD.

**Keywords:** Incretin agonists, GLP-1 receptor agonists, tirzepatide, semaglutide, multi-omics, plasma proteomics, metabolomics, inflammation, biological aging, metabolic aging clock, weight loss, pharmacodynamics, mitochondria

## Abstract

The benefits of incretin agonists, including glucagon-like peptide-1 receptor agonists and dual GIP/GLP-1 receptor agonists, are largely attributed to glycemic control and weight loss, as these therapies enhance insulin secretion, suppress glucagon release and delay gastric emptying to produce satiety. Yet treatment is also associated with lower risk of cardiometabolic, renal and other chronic diseases. Whether these broader benefits depend on weight loss itself or reflect direct effects of treatment on physiology remains unclear, as does the basis of the marked variation in how patients respond. Resolving these questions has been difficult because weight loss and drug effects occur together, and because molecular studies of treated patients have been limited in scope. Here we profiled paired plasma samples collected before and after treatment from 165 individuals drawn from DynamiQ, a biorepository that links plasma samples to longitudinal clinical records, combining mass spectrometry-based proteomics, a targeted inflammation proteomics panel (NULISAseq) and non-targeted metabolomics to capture metabolic function, inflammation and markers of chronic disease. Treatment was associated with broad remodeling of the plasma proteome and metabolome, but the extent to which this remodeling depended on weight loss differed across the molecular layers. The metabolome carried a lipid signature that scaled with the weight lost, whereas proteomic and inflammation-panel changes largely did not depend on weight loss. Among these changes, treatment was associated with lower circulating mitochondrial oxidative proteins and cardiolipins, consistent with relief of lipid-driven metabolic stress, and with a shift of the inflammation panel toward a less inflamed state. A metabolic aging clock showed that age acceleration decreased by a mean of 2.96 years after treatment (95% confidence interval −4.95 to −0.98; P = 3.7 × 10⁻³), a change that did not depend on weight loss or on the interval between samples, with the largest reductions in participants starting from the highest age acceleration, most of whom had diabetes. Treatment-associated features overlapped with features associated with cardiometabolic, renal and hepatic disease. Before treatment, higher systemic inflammation, reflecting chronic tissue stress and repair, was associated with less subsequent weight loss. Together, these findings indicate that incretin agonist treatment is associated with lower markers of metabolic and inflammatory stress and with a younger metabolic age, changes that do not depend on weight loss, identify baseline inflammation as an axis of heterogeneity in treatment response, and suggest that biorepositories linking longitudinal multi-omic profiling to clinical records can relate drug pharmacodynamics to disease risk and to variation in patient response.

**Synopsis:** Paired multi-omics in 165 individuals before and after incretin agonist treatment separates a triacylglycerol profile that correlates with weight loss from a broader response not dependent on it, accompanied by a younger metabolic age.

- Most proteomic and inflammation-panel changes after incretin agonist treatment are not dependent on weight loss, whereas the metabolome carries a triacylglycerol signature that scales with weight loss.
- Treatment is associated with lower circulating mitochondrial oxidative proteins and cardiolipins without signs of tissue injury, consistent with relief of lipid-driven metabolic stress.
- Metabolic age acceleration decreases by about three years, a change not dependent on weight loss, and treatment-associated features overlap with features of cardiometabolic, renal and hepatic disease risk.
- Higher systemic inflammation before treatment, reflecting chronic tissue stress and repair, is associated with less subsequent weight loss.

## Introduction

Incretin agonists, including glucagon-like peptide-1 receptor agonists (GLP-1RAs) and the dual GIP/GLP-1 receptor agonist tirzepatide, have transformed the treatment of type 2 diabetes and obesity, producing weight reductions of 15–20% in clinical trials and reshaping both clinical practice and public health discourse around metabolic disease^1,2^. Beyond glycemic control and weight loss, a growing body of evidence links their use to reduced risk across a broad range of conditions, including cardiovascular events^3–6^, Alzheimer’s disease^7,8^ and substance use disorders^8^, as well as reduced alcohol consumption in alcohol use disorder^9^ and improvement in metabolic dysfunction-associated steatohepatitis (MASH)^10,11^. This breadth of response raises the question of which other physiological processes are influenced by treatment, and whether the beneficial effects are, at least in part, not dependent on the degree of weight loss achieved.

Preclinical work has begun to characterize systemic effects of these drugs that do not depend on weight loss. In aged mice, GLP-1RA therapy counteracts age-related changes across multiple organs and molecular layers at a dose with minimal effect on food intake or body weight, and these effects depend largely on hypothalamic GLP-1 receptor signaling^12^. More recently, late-life semaglutide treatment was shown to slow aging and extend lifespan in female mice^13^. Direct human evidence linking treatment to biological aging has also begun to emerge: in a randomized trial in people with HIV-associated lipohypertrophy, semaglutide slowed epigenetic aging^14^, and in a single-arm pilot study in people with HIV and metabolic dysfunction-associated steatotic liver disease, changes in epigenetic clocks correlated with the clinical response to semaglutide^15^. GLP-1RAs also have anti-inflammatory and immunomodulatory effects, and in a small human study circulating inflammatory markers fell independently of changes in body weight and glycemia^16^. Together, these findings raise a central open question: which of the clinical benefits attributed to incretin agonist therapy reflect weight loss itself, and which reflect effects of the drug that do not depend on weight loss?

Progress on this question has been limited by the scope of available human evidence. The largest proteomic study of treatment in humans, using serum from the STEP 1 and STEP 2 semaglutide trials, identified broad changes spanning body weight regulation, glycemic control, lipid metabolism and inflammation, and found that a subset persisted after statistically accounting for weight loss and glycemic improvement^17^. However, this and other prior work has examined a single molecular layer, in controlled trial populations that may not reflect real-world treatment, and has not linked biomarker changes to biological age or to real-world disease outcomes. Similarly, a large study using electronic health record (EHR) data has documented substantial heterogeneity in real-world weight loss, with some individuals losing more than 15% of body weight and others losing little or none on the same agents. This study identified clinical and demographic correlates of response, but without accompanying molecular data^18^. Resolving these questions requires molecular measurements across several layers in the same people before and after treatment, together with a means of relating the changes to disease risk. Clinical trials are poorly suited to this: molecular profiling within trials is costly and usually limited to a single layer, and trials cannot sample participants years before a diagnosis or before and after treatment begun in routine care. Biorepositories that profile repeated samples from large numbers of individuals on a single platform, linked to their clinical records, offer a complementary route.

Here we address these gaps in a real-world cohort of 165 participants with paired plasma samples collected before and after incretin agonist treatment. By combining mass spectrometry-based proteomics, a targeted inflammation proteomics panel and non-targeted metabolomics, we measure the response to treatment across complementary layers of physiology in the same individuals. We show that treatment is associated with broad remodeling of the plasma proteome and metabolome, and that the extent to which this remodeling depends on weight loss differs between molecular layers. The metabolome carries a lipid signature, dominated by triacylglycerols, that scales with each participant’s weight loss. By contrast, the proteomic and inflammation-panel changes largely do not depend on weight loss. These include a reduction in circulating mitochondrial oxidative proteins and a shift of the inflammation panel toward a less inflamed state, driven by lower levels of chemokines, T-cell costimulatory molecules and myeloid growth factors. Treatment is also associated with reduced metabolic age acceleration that does not depend on weight loss, and the size of this reduction tended to be larger in individuals with stronger mTORC1 and PI3K/AKT growth signaling and immune responses. Because the cohort was drawn from a biorepository that samples individuals before diagnosis, we could also relate the treatment response to real-world disease risk, finding overlap with features associated with the cardiometabolic, renal and hepatic diseases these drugs are known to prevent. Finally, we find that higher systemic inflammation before treatment, reflecting chronic tissue stress and repair, is associated with less subsequent weight loss. Together, these findings suggest that the benefits of incretin agonists extend beyond weight loss to the metabolic and inflammatory biology of aging, and that a patient’s molecular state before treatment may bias treatment outcomes.

## Results

### Cohort and systemic multi-omic response to treatment

We sought to understand the molecular impact of incretin treatment, and whether part of the response does not depend on weight loss, through mass spectrometry-based proteomics, a targeted inflammation proteomics panel and metabolomics measured in plasma, an easily sampled specimen that reflects systemic physiology^19^. We identified 165 participants (330 paired pre-and post-treatment plasma samples) treated with a GLP-1-based therapy (semaglutide, liraglutide, dulaglutide or tirzepatide) for at least 90 days (Supplementary Fig. S1; Supplementary Table 1). This cohort was curated from DynamiQ biorepository (Sapient Bioanalytics) linked with multi-omics and real-world data, comprising 65,847 total plasma samples collected longitudinally from 11,920 individuals (Fig. 1c). To stratify the heterogeneous weight loss response, percent weight change from baseline classified participants into three response groups whose trajectories remained distinct through 18 months of follow-up: weight loss of 5% or more (n = 103), weight loss of 0–5% (n = 22) and no weight loss (n = 31); weight data were insufficient to classify the remaining 9 participants (Fig. 1a,b). By integrating proteomics and metabolomics data, we sought to systematically characterize tens of thousands of molecular features that together define the functional chemistry of the human body, capturing signals of cell death, growth and immune response alongside the metabolic pathways that sustain physiological function.

**Figure 1.**
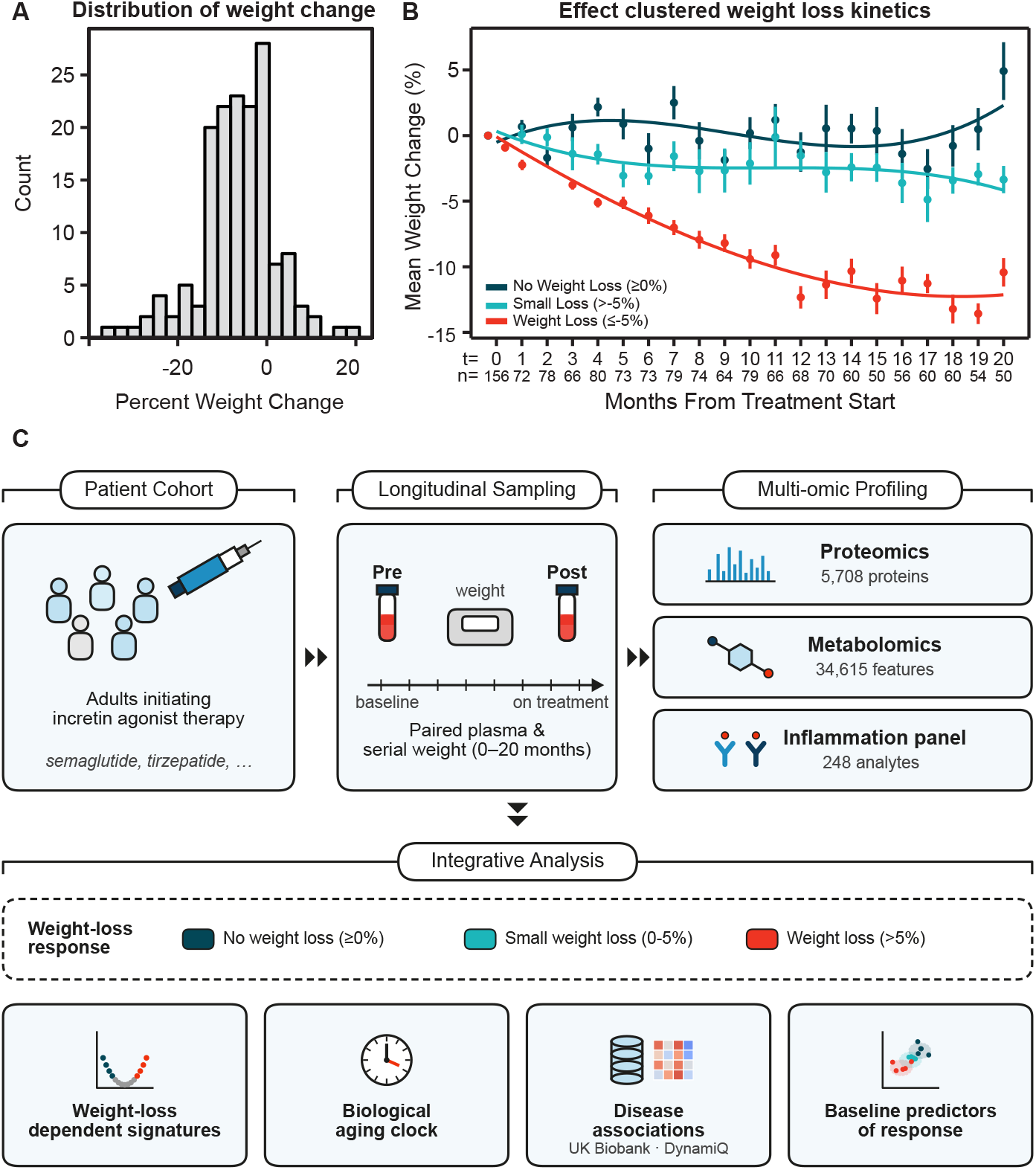
Cohort and weight-loss characterization. a, Histogram of percent weight change from baseline (n = 156 participants with evaluable weight data). b, Mean weight change over time from treatment start by response category: no weight loss (weight change ≥ 0%, n = 31), weight loss of 0–5% (n = 22) and weight loss of 5% or more (n = 103); points show group means at each timepoint, error bars show 95% confidence intervals and curves show a smoothed fit within each group. c, Study design: selection of 165 participants (330 paired samples) treated with an incretin agonist from the DynamiQ biorepository, paired pre-and post-treatment sampling, multi-omic profiling (proteomics, metabolomics and inflammation panel) and the integrative analyses performed.

We took a global view of the features that were altered comparing post-treatment (post) plasma to paired samples pre-treatment (pre), which identified widespread molecular changes after treatment at a false discovery rate (FDR) below 0.1 (Fig. 2a; Supplementary Tables 2–5). These included 1,080 of 5,708 proteins (444 decreased, 636 increased), 5,621 of 34,615 metabolite features (3,845 decreased, 1,776 increased) and 35 of 248 inflammation panel analytes (29 decreased, 6 increased), representing remodeling of plasma composition. To ask what the changed proteins and metabolites have in common, we tested for enrichment of pathways and metabolite classes among the increased and decreased features. Among increased proteins, pathway enrichment identified mTORC1 signaling (52↑, 11↓; FDR = 2.7 × 10⁻⁴), Myc targets (43↑, 10↓; FDR = 2.1 × 10⁻⁵), glycolysis (28↑, 11↓; FDR = 9.0 × 10⁻³) and PI3K/AKT/mTOR signaling (24↑, 2↓; FDR = 0.019) (Fig. 2b). All twelve detected mTOR-axis regulators increased after treatment, while no canonical readout of mTORC1 activity changed significantly (Fig. 2d). The activators included the direct mTORC1 activator RHEB (+0.17 log2 fold change, FDR = 8.9 × 10⁻¹⁰), its upstream kinase AKT1 (+0.12, FDR = 4.5 × 10⁻⁴), the mTOR assembly factor TELO2 (+0.31, FDR = 0.023) and the PKA catalytic subunits PRKACA (+0.23, FDR = 1.3 × 10⁻⁸) and PRKACB (+0.11, FDR = 1.1 × 10⁻⁶) (Fig. 2d; Supplementary Table 6). Because GLP-1 receptor signaling can activate mTORC1 through PKA^20^, a direct drug effect is plausible. However, the negative regulators PTEN (+0.25, FDR = 0.034) and STK11/LKB1 (+0.23, FDR = 0.025) also increased (Fig. 2d). These increases may reflect an adaptive response to weight loss, a direct effect of treatment, or both.

**Figure 2.**
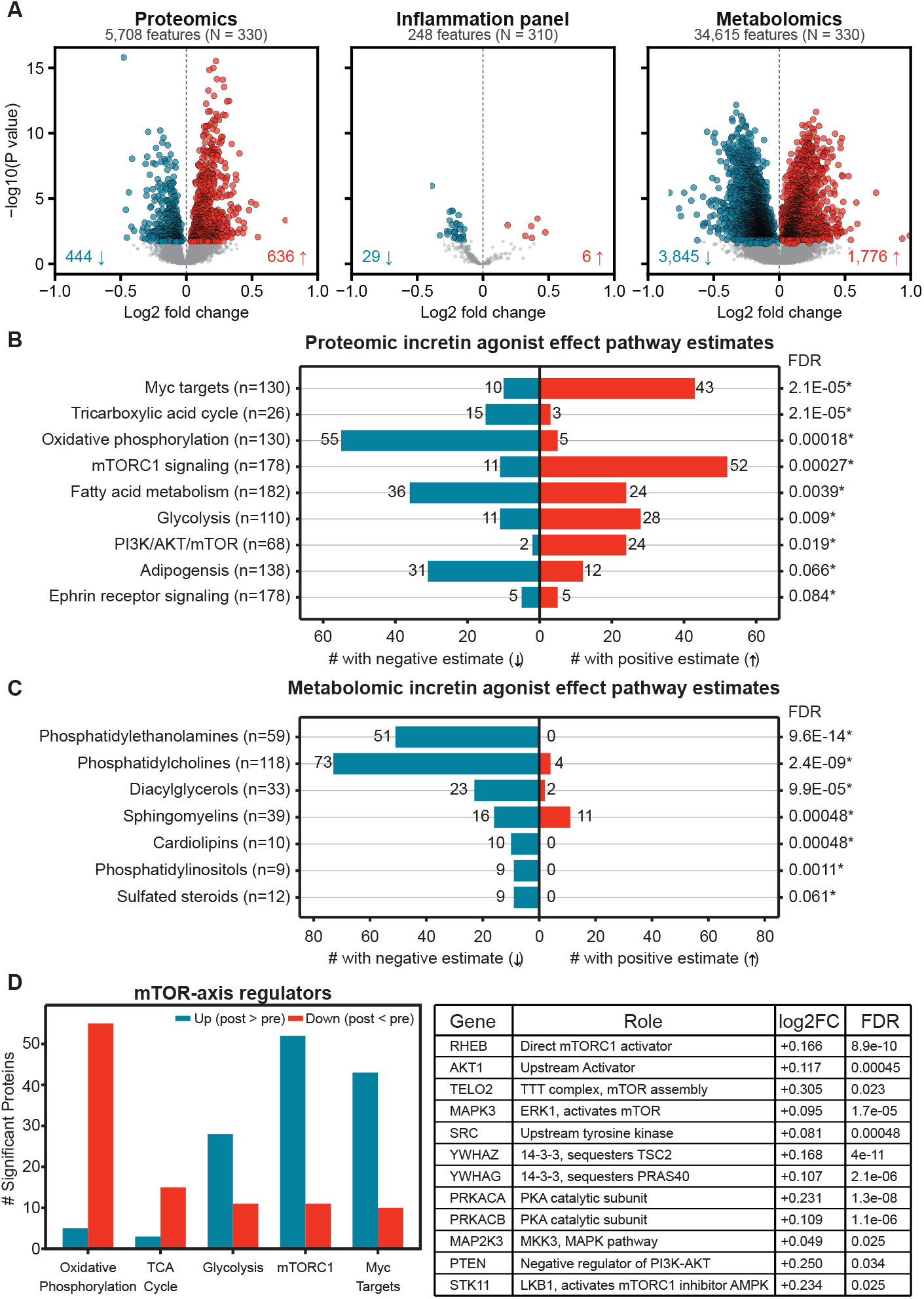
Systemic multi-omic response to incretin agonist treatment. a, Paired pre-and post-treatment differential abundance for proteomics (left), the inflammation panel (middle) and metabolomics (right). Blue points, significantly decreased; red points, significantly increased (two-sided paired t-test, FDR < 0.1); gray points, not significant. Numbers of significantly decreased and increased features and the total number tested are shown in each panel; n = 330 samples from 165 participants for proteomics and metabolomics, 310 samples from 155 participants for the inflammation panel. b, Enrichment of protein gene sets among significantly changed proteins; bars show the number of members with negative (blue, left) or positive (red, right) effect estimates; FDR from one-sided Fisher’s exact test at right; asterisks denote FDR < 0.1. c, As in b, for metabolite compound classes. d, Direction of change for detected components of the mTOR axis, grouped into pathway activators, pathway inhibitors and canonical activity readouts; points show log2 fold change with 95% confidence intervals; filled points denote FDR < 0.1. Activity readouts that were detected but unchanged are shown to support the statement that these data do not demonstrate directional mTOR signaling.

Many of the decreased proteins were related to mitochondrial function. Oxidative phosphorylation (5↑, 55↓; FDR = 1.8 × 10⁻⁴), the tricarboxylic acid (TCA) cycle (3↑, 15↓; FDR = 2.1 × 10⁻⁵) and fatty acid metabolism (24↑, 36↓; FDR = 3.9 × 10⁻³) were enriched among them (Fig. 2b). Adipogenesis showed the same tendency (12↑, 31↓; FDR = 0.066). The decreased oxidative phosphorylation proteins spanned complex I (NDUFS1, NDUFS3, NDUFA10), complex II (SDHA, SDHB), complex III (UQCRC2), complex IV (COX5A, COX5B) and ATP synthase (ATP5F1A, ATP5PO). The decreased TCA cycle enzymes included citrate synthase (CS), aconitase (ACO2), isocitrate dehydrogenase (IDH3A) and malate dehydrogenase (MDH2). The fatty acid metabolism set included the mitochondrial β-oxidation enzymes CPT2, ACADVL, ACADM, HADHA and HADHB (Supplementary Fig. S2a). Together, these classes of core mitochondrial metabolism proteins point to an altered state of the mitochondria, which was further supported by the metabolomics data. All detected cardiolipins, the signature phospholipid of the inner mitochondrial membrane, decreased (0↑, 10↓; FDR = 4.8 × 10⁻⁴). Phosphatidylethanolamines (0↑, 51↓; FDR = 9.6 × 10⁻¹⁴), phosphatidylcholines (4↑, 73↓; FDR = 2.4 × 10⁻⁹), diacylglycerols (2↑, 23↓; FDR = 9.9 × 10⁻⁵), phosphatidylinositols (0↑, 9↓; FDR = 1.1 × 10⁻³) and sulfated steroids (0↑, 9↓; FDR = 0.061) also decreased. These decreases are consistent with the reductions in diacylglycerols and triglyceride-rich lipoproteins reported with incretin agonists^21,22^. Sphingomyelins were the only class with a substantial number of increased species, and they changed in both directions (11↑, 16↓; FDR = 4.8 × 10⁻⁴) (Fig. 2c).

A treatment-associated reduction in the circulating mitochondrial signature could arise in two ways: mitochondrial metabolism in source tissues could have changed, or less mitochondrial material could be released into the circulation from stressed or dying cells^23,24^. Mitochondrial contents released from injured cells, including formyl peptides and mitochondrial DNA, act as damage-associated molecular patterns that activate neutrophils through formyl peptide receptor 1 and Toll-like receptor 9 and can drive systemic inflammation^25^. A change in cell death should therefore engage the receptors that sense tissue damage and the inflammatory cytokines downstream of them. Yet most injury-responsive markers on the inflammation panel were not dysregulated: of 43 analytes spanning inflammatory cytokines that respond to cell and tissue death, receptors that sense damage-associated molecular patterns, death ligands and their receptors, macrophage and clearance markers and stress hormones, 40 were unchanged or lower after treatment, including the canonical injury cytokines IL-1β, IL-6, TNF, CXCL8, IL-18 and CRP (Supplementary Fig. S2b). Moreover, markers of macrophage activity and apoptotic-cell clearance were unchanged or lower, including CSF1R (−0.26, FDR = 0.052) and CD93 (−0.21, FDR = 0.039). FGF21, a hormone induced by mitochondrial stress^26^, decreased (−0.31, FDR = 0.054), whereas GDF15, a stress-induced cytokine that also rises with mitochondrial dysfunction^27^, did not change (FDR = 0.80). More broadly, the inflammation panel shifted toward a less inflamed state after treatment, with 29 of the 35 significantly changed analytes decreasing (Fig. 2a). The decreases spanned chemokines that recruit and position immune cells (CCL25, CCL19, CCL21, CCL13, CCL16, CCL1), costimulatory molecules and cytokines that drive T-cell and antigen-presenting-cell activation (TNFRSF4/OX40, TNFRSF18/GITR, ICOSLG, CD83, IL-12B), and growth factors for myeloid cells (CSF2, FLT3LG, KITLG). Together, these changes indicate a coordinated reduction in immune activation and recruitment rather than a response to tissue injury.

Lower circulating oxidative proteins do not necessarily indicate lower oxidative capacity. In obese, insulin-resistant people, maximal hepatic mitochondrial respiration is 4.3-to 5.0-fold higher than in lean people, an adaptation to excess lipid supply^28^. In obese rats with steatotic liver disease, semaglutide combined with PYY3-36 lowered hepatic oxidative phosphorylation gene expression, mitochondrial respiration and mitochondrial H₂O₂ production and reduced hepatocellular stress, whereas food restriction and weight loss had no significant effect on mitochondrial respiration or oxidative phosphorylation genes^29^. Fat loss itself requires sustained oxidation of stored lipids. The concurrent decreases in cardiolipins, phospholipids and diacylglycerols (Fig. 2c) are consistent with lower lipid flux through the mitochondria rather than with a loss of mitochondrial function. Reduced release of mitochondrial material, through lower cellular stress or improved mitochondrial quality control, may also contribute to this observed decrease in plasma proteins. Nonetheless, the absence of an injury or inflammatory signal may support metabolic reprogramming toward a lower lipid-driven oxidative load over mitochondrial loss or failure. Such a reduction in lipid load could arise in two ways: as a downstream consequence of fat loss itself, or as a direct effect of treatment that does not require weight loss.

### Molecular responses that do and do not depend on weight loss

Incretin agonists cause weight loss, so any molecular change after treatment could reflect weight loss itself or an effect of the drug that does not depend on it. To separate these, we correlated each feature’s change after treatment with percent weight change. Weight loss explained a defined part of the metabolomic response but almost none of the proteomic or inflammatory response (Fig. 3a). Only 1 of 5,708 proteins and 14 of 248 inflammation panel analytes (11 negatively, 3 positively) correlated with percent weight change. The increase in mTOR-axis proteins, the decrease in mitochondrial oxidative proteins and the broad reduction in inflammatory analytes therefore did not depend on the extent of weight loss and are more likely to reflect incretin agonist treatment than the weight loss it produces. By contrast, and as expected, the metabolome carried a measurable signature of weight loss: 596 of 34,615 metabolite features correlated with percent weight change (533 positively, 63 negatively). The weight-loss-dependent metabolites were dominated by triacylglycerols. All 59 significant triacylglycerols correlated positively with weight change, so that greater weight loss accompanied larger decreases (FDR = 2.7 × 10⁻¹⁹). Very-long-chain dicarboxylic acids, products of fatty acid ω-oxidation, followed the same pattern (0 negative, 7 positive; FDR = 5.0 × 10⁻³). Sphingomyelins changed in both directions with treatment (Fig. 2c) and also split in their relationship to weight loss (9 negative, 4 positive; FDR = 8.9 × 10⁻³) (Fig. 3b). Triacylglycerols are the storage form of fat, so their correlation with weight loss likely reflects the loss of fat mass.

**Figure 3.**
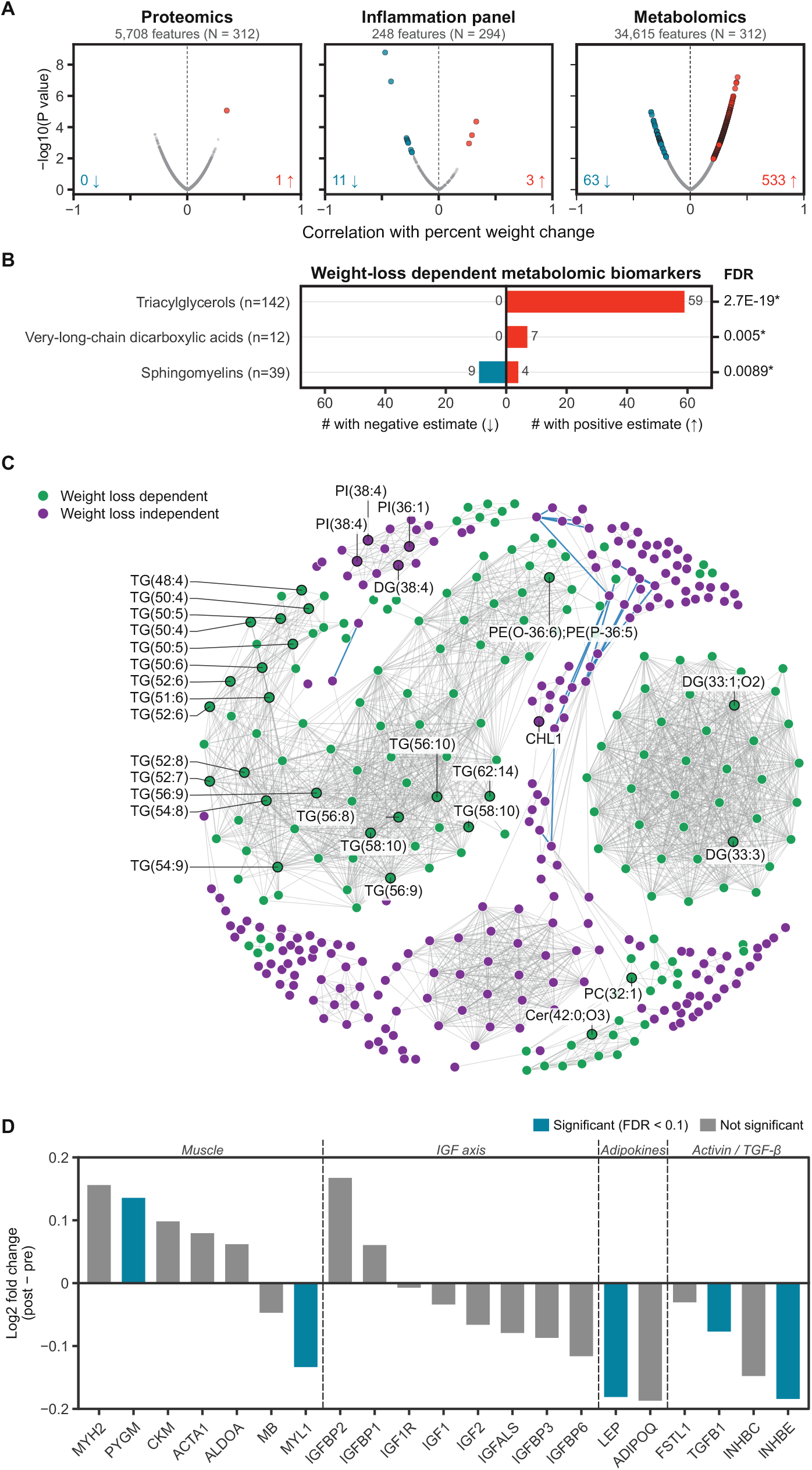
Dissociation of weight loss from the molecular response. a, Regression between biomarker fold change and percent weight loss, by omics layer, formatted as in Fig. 2a; n = 312 samples from 156 participants for proteomics and metabolomics, 294 samples from 147 participants for the inflammation panel. b, Enrichment of metabolite compound classes among features whose fold change is associated with percent weight loss, formatted as in Fig. 2c. c, Spearman correlation network among significant pharmacodynamic biomarkers, restricted to edges with absolute r > 0.7; node color indicates whether a biomarker’s change is dependent (green) or not dependent (purple) on weight loss, as defined in Methods. d, Log2 fold change with 95% confidence intervals for detected members of the activin/TGF-beta axis, the IGF axis, adipokines and muscle structural and metabolic proteins; filled points denote FDR < 0.1; axis members not detected in plasma are listed beneath the panel rather than plotted.

In contrast to these storage lipids, the membrane lipids that changed with treatment did not correlate with weight loss. Cardiolipins, phosphatidylethanolamines and phosphatidylcholines were not among the weight-loss-dependent classes, extending the response that does not depend on weight loss from mitochondrial proteins to membrane lipids, including the mitochondrial lipid cardiolipin. To see how the components that do and do not depend on weight loss relate, we built a correlation network of the significant features (Fig. 3c). The two sets of features separated into distinct clusters. Triacylglycerols formed the core of the weight-loss-dependent clusters. The clusters not dependent on weight loss contained phosphatidylinositols and the adhesion molecule CHL1, together with treatment-responsive proteins spanning the mitochondrial respiratory chain, complement and coagulation, cell adhesion and lipid transport (Supplementary Table 20). The treatment response therefore comprises two distinguishable molecular profiles: a lipid profile that scales with weight loss, and a broader proteomic and metabolic profile that does not depend on it.

### Incretin treatment remodels the tissue-mass axes reflected in the plasma

Loss of lean mass alongside fat is a recognized feature of weight loss on incretin agonists^1,30^, and the circulating proteins that regulate muscle and adipose mass offer a view of how these tissues respond to treatment. We therefore examined the muscle, insulin-like growth factor (IGF), adipokine and activin/TGF-β axes (Fig. 3d; Supplementary Table 7), the pathways that set how much tissue the body builds and stores. IGF signaling drives muscle protein synthesis and adipokines report on fat mass, whereas activin signaling restrains muscle growth, and blocking activin receptors largely preserves lean mass during semaglutide treatment^30^. Leptin decreased (−0.18, FDR = 0.045), consistent with reduced fat mass, and within the activin/TGF-β axis both INHBE (−0.18, FDR = 0.012) and TGFB1 (−0.08, FDR = 3.1 × 10⁻³) decreased. The IGF axis remained stable, with no significant change in IGF1, IGF2, IGF1R or the IGF-binding proteins. Because circulating IGF-1 decreases with sustained energy and protein restriction^31^, its stability indicates that the anabolic growth axis was maintained despite reduced intake. Among muscle proteins, the muscle glycogen phosphorylase PYGM increased (+0.14, FDR = 1.1 × 10⁻⁴) and the fast-twitch myosin light chain MYL1 decreased (−0.13, FDR = 3.2 × 10⁻⁴). Meanwhile, the established circulating markers of muscle injury, creatine kinase (CKM) and myoglobin (MB), remained unaffected along with ACTA1 and MYH2, a pattern that could point to selective remodeling of muscle glycogen metabolism and fiber composition rather than muscle breakdown.

Of these changes, the decrease in INHBE is notable because genetic loss of INHBE is associated with favorable metabolic health. INHBE encodes activin E, a liver-derived member of the activin family whose expression is largely confined to hepatocytes^32^. In human genetic studies, heterozygous loss-of-function variants in INHBE are associated with a healthier distribution of body fat^32,33^, together with a more favorable metabolic profile and roughly 28% lower odds of type 2 diabetes^32^, and on the strength of this evidence an siRNA that silences hepatic INHBE has entered clinical testing for obesity^34^. INHBE was also the only protein in the plasma proteome whose change scaled with the extent of weight loss (Fig. 3a). Incretin agonist treatment thus lowers a liver-derived hormone whose genetic loss protects against metabolic disease, and it does so in proportion to the weight lost. This places a genetically validated therapeutic target directly within the weight-loss-dependent profile of the treatment response, alongside the triacylglycerols that correlated with weight loss in the metabolome. Taken together, these observations indicate that incretin agonist treatment moves plasma composition toward a profile associated with metabolic health, raising the question of whether it also shifts the circulating proteome toward a biologically “healthier” state.

### Incretin agonist treatment is associated with reduced biological age acceleration

Biological age, as opposed to chronological age, can be impacted by lifestyle factors, disease or pathology, and genetic factors such that two individuals of identical chronological age can differ biologically^35,36^. The shift of plasma toward a profile associated with metabolic health led us to ask whether incretin agonist treatment also slows, or even reverses, biological aging. To infer this, we used a metabolic aging clock: a model trained on healthy people to predict chronological age from the levels of a small set of plasma metabolites that change steadily with age. When applied to a new sample, the clock returns a predicted biological age. The gap between that prediction and the person’s chronological age, termed age acceleration (also termed Δage) indicates whether their metabolism resembles that of someone older or younger than they are (Fig. 4a). We previously developed such a clock on this platform^36^ trained on plasma metabolite profiles from 887 healthy adults selected from the DynamiQ biorepository to span six decades of age, using robust age-associated features in an interpretable linear model. Applied to people with chronic disease, it reported biological ages several years older than their chronological ages^36^. In individuals who underwent kidney transplantation, that excess fell sharply after surgery, suggesting that the clock captures systemic aging that responds to treatment^36^. We retrained the clock for the present analysis on 183 healthy control participants measured on the same platform as the treated cohort, retaining 20 of the 32 candidate features and detrending predictions against chronological age to correct the compression of predicted age toward the cohort mean (Methods). The retrained clock predicted chronological age closely across an age range of 22.5 to 83.3 years, both in the training set (190 samples from 129 participants; Pearson r = 0.88, R² = 0.70, median absolute error 7.01 years) and in a held-out set (92 samples from 54 participants; r = 0.87, R² = 0.62, median absolute error 7.11 years) (Fig. 4b,c; Supplementary Table 8). Applied to the treated cohort, the clock placed participants well above their chronological age before treatment, with a mean age acceleration of about +10 years and a correspondingly looser fit to chronological age (r = 0.47; Supplementary Fig. S3b). This is consistent with a population enriched for diabetes and obesity, conditions the clock was previously shown to accelerate biological age^36^.

**Figure 4.**
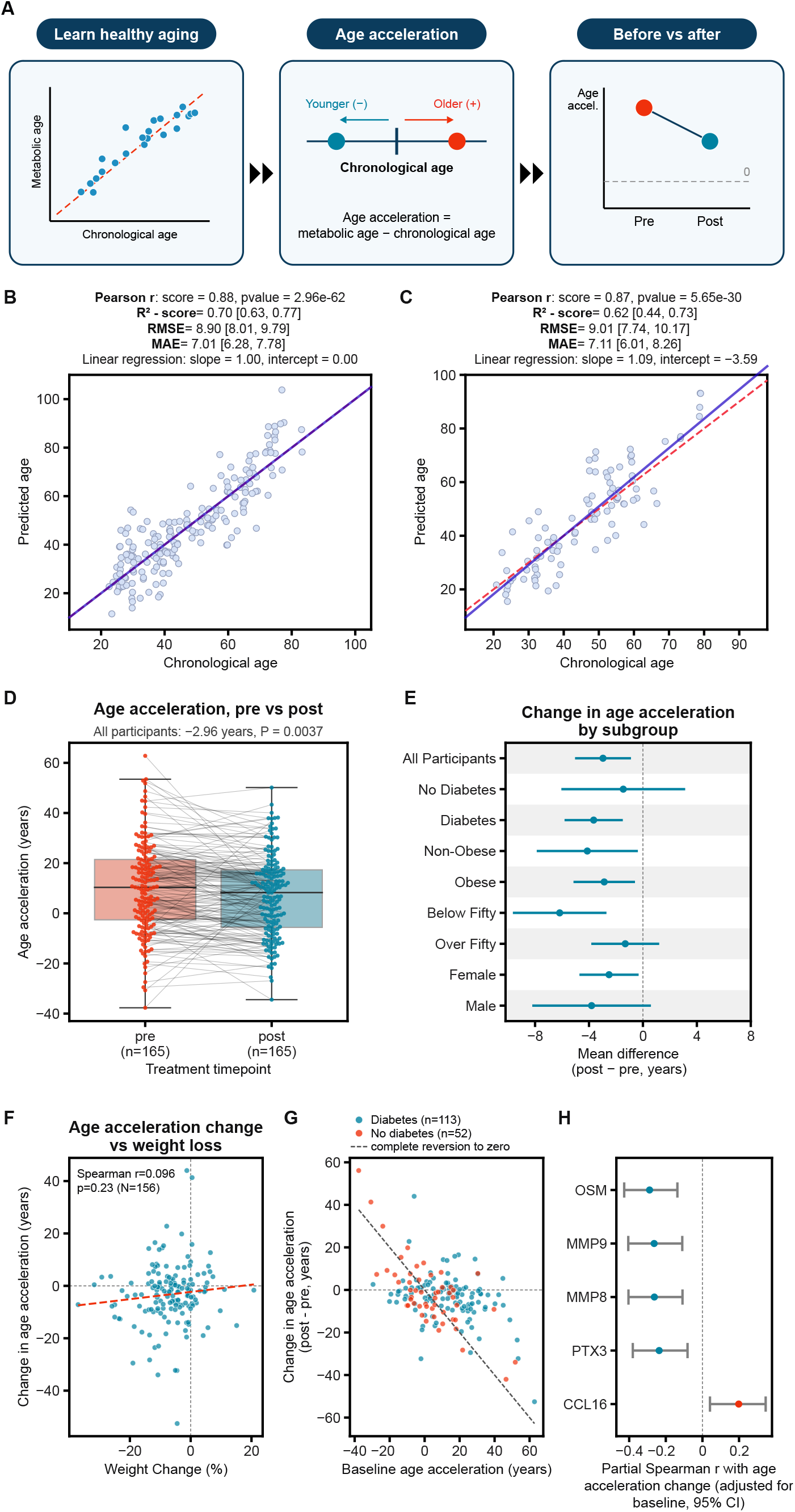
A metabolic aging clock shows reduced biological age acceleration during incretin agonist treatment. a, Analysis design. A clock trained on healthy individuals predicts age from plasma metabolites; age acceleration is predicted (metabolic) age minus chronological age, positive when a person’s metabolism resembles that of someone older; age acceleration was then compared between each participant’s pre-and post-treatment samples. b, Detrended predicted age versus chronological age in the clock training set (190 samples from 129 participants); solid line, linear fit; dashed line, identity. c, As in b, for the held-out validation set (92 samples from 54 participants not used in training). Pearson r = 0.88 and 0.87, R2 = 0.70 and 0.62, median absolute error 7.01 and 7.11 years respectively. d, Age acceleration in paired pre-and post-treatment samples (n = 165 participants); box shows median and interquartile range; whiskers show 1.5x the interquartile range; gray lines connect paired samples from the same individual. e, Forest plot of the mean within-person change in age acceleration (post minus pre) for all participants and within demographic and clinical strata; points show the mean difference, horizontal bars the 95% confidence interval, dashed line no change. Strata are within-group tests; between-group contrasts are reported in Supplementary Table 15. f, Within-person change in age acceleration versus percent weight loss (n = 156; Spearman r = 0.096, P = 0.23); dashed line, linear fit. g, Within-person change versus baseline age acceleration, colored by diabetes status (diabetes, n = 113; no diabetes, n = 52); the dashed line marks complete reversion of baseline age acceleration to zero. h, Partial Spearman correlation, adjusted for baseline age acceleration, between the within-person change in age acceleration and the treatment-associated change in each inflammation-panel analyte associated at FDR < 0.1 (OSM, MMP9, MMP8, PTX3 and CCL16); bars show 95% confidence intervals.

We compared age acceleration between each participant’s paired pre-and post-treatment samples and found that it decreased after treatment by a mean of 2.96 years (95% confidence interval −4.95 to −0.98; paired t-test P = 3.7 × 10⁻³, n = 165) (Fig. 4d; Supplementary Table 9). We assessed the impact of several possible confounders. This decrease in age acceleration was not a function of time: it was unrelated to the interval between paired samples, the duration of treatment, the calendar date of the baseline sample (Supplementary Table 10), erythrocyte contamination (Supplementary Note 1) and sample storage time. The decrease in age acceleration did not scale with the extent of weight loss (Spearman r = 0.096, P = 0.23, n = 156) (Fig. 4f), which places the slowing of metabolic aging alongside the proteomic and inflammatory changes among the treatment-associated changes that do not depend on weight loss. Because participants entered treatment with a mean age acceleration of about +10 years, a decrease of nearly three years corresponds to almost a third of that excess. The decrease also varied considerably from one participant to the next, which gave us the opportunity to ask which components of the broader molecular response were associated with the decrease, and therefore which parts of the treatment response are most closely tied to a younger metabolic state.

To identify these components, we scored each participant’s response in the pathways that defined the proteomic changes in Fig. 2b and in related immune gene sets. Each score was the mean z-scored change of the pathway’s treatment-responsive proteins, so a higher score means that a participant’s proteins in that pathway rose more with treatment. We correlated each score with the change in age acceleration after adjusting for baseline age acceleration. The pathways that increased with treatment were all negatively associated with the change, meaning that participants whose growth-signaling and immune proteins rose most showed the largest reductions in metabolic age. These associations trended for mTORC1 signaling (partial Spearman r = −0.212, FDR = 0.064), PI3K/AKT/mTOR signaling (r = −0.188, FDR = 0.064), innate immune response (r = −0.187, FDR = 0.064), complement (r = −0.180, FDR = 0.064), Myc targets (r = −0.170, FDR = 0.070) and inflammatory response (r = −0.165, FDR = 0.070), with glycolysis and coagulation trending in the same direction (Supplementary Fig. S4b; Supplementary Table 12). In contrast, the mitochondrial and lipid pathways that decreased with treatment showed no association, including oxidative phosphorylation (r = +0.088), the TCA cycle (r = +0.052), adipogenesis (r = +0.029) and fatty acid metabolism (r = +0.001). The mitochondrial reprogramming described above thus appears to be a shared feature of the response to incretin agonists, whereas the strength of the growth-signaling and immune response varied between participants and correlated with the change in age acceleration.

Because the metabolic clock is derived from metabolites, we examined whether the proteomic response also intersected an independent, protein-based measure of biological age. Three proteins from a proteomic aging clock developed in the UK Biobank^37^ decreased with treatment: LECT2 (log2 fold change −0.26, FDR = 0.029) and endoglin (ENG; log2 fold change −0.066, FDR = 0.067) from its 20-protein core, and tenascin-X (TNXB; log2 fold change −0.13, FDR = 0.068). Their concordant decrease is consistent with incretin agonist treatment lowering proteins that independently mark biological age, extending the link between treatment and aging to a second molecular layer and to an aging signature derived from more than 45,000 individuals. LECT2 is a hepatokine that links obesity to skeletal muscle insulin resistance^38^, and its decrease adds to the liver-derived signals, including INHBE, that were lowered by treatment.

The metabolic clock and proteomic data thus converge on a younger systemic state, and the pathway analysis indicates that the reduction in metabolic age was greatest in participants with the strongest immune and growth-signaling response. We sought direct evidence for this immune component in the inflammation panel, which measures individual immune mediators. After adjustment for baseline age acceleration, five analytes were associated with the change in age acceleration: oncostatin M (OSM; partial Spearman r = −0.289, FDR = 9.9 × 10⁻³), MMP9 (r = −0.264, FDR = 0.012), MMP8 (r = −0.262, FDR = 0.012), pentraxin 3 (PTX3; r = −0.236, FDR = 0.028) and the liver-derived chemokine CCL16 (r = +0.198, FDR = 0.096) (Fig. 4h; Supplementary Table 14). The first four were among the few inflammation-panel analytes that rose with treatment (Fig. 2a), and CCL16 fell. In every case, a larger treatment-associated change accompanied a larger reduction in age acceleration, linking the clock response to mediators of tissue remodeling and innate immunity.

Having identified which molecular programs were linked with the change in metabolic age, we next asked who benefited most from incretin treatment. The decrease in age acceleration was consistent in direction across demographic and clinical strata (Fig. 4e; Supplementary Fig. S3c–e). It was significant in participants with diabetes (−3.65 years, P = 6.8 × 10⁻⁴, n = 113), in those below fifty years of age (−6.17 years, P = 5.7 × 10⁻⁴, n = 56), in women (−2.51 years, P = 0.020, n = 108), and in both obese (−2.87 years, P = 0.011, n = 108) and non-obese participants (−4.13 years, P = 0.028, n = 46). In men, the decrease was larger than in women (−3.81 years, n = 56) but did not reach significance (P = 0.083) in this smaller stratum. The largest decreases occurred in participants who started with the highest age acceleration (Fig. 4g), and participants with diabetes carried most of this excess at baseline (median +13.7 versus +1.0 years without diabetes; Supplementary Table 15). Participants with diabetes contributed substantially to the incretin-associated molecular signal. Within this stratum (n = 113), 5,461 metabolite features, 1,090 proteins and 34 inflammation panel analytes changed with treatment, compared with none, none and one, respectively, in the smaller stratum of participants without diabetes (n = 52), and within the diabetes stratum 825 metabolite features correlated with weight loss, compared with 596 in the full cohort (Fig. 3a) (Supplementary Table 2; Supplementary Figs S5b, S6, S7 and S8). Taken together, the response to incretin agonists was largest in those with the greatest metabolic burden, although no between-group contrast remained significant after adjustment for baseline age acceleration (Supplementary Table 15), so the stratum differences describe where the effect was detectable rather than a demonstrated difference in treatment response. Accelerated metabolic aging is itself a feature of chronic disease^36^, and the participants who responded most were those whose metabolic burden placed them at greatest risk. We therefore next asked whether the molecular features that respond to incretin agonists also overlap with features associated with the risk of developing disease.

### Treatment-associated features overlap with features of disease risk

Incretin agonists reduce the incidence of cardiovascular, renal and metabolic disease in clinical trials^3–6,10,11^ and real-world cohorts^8^, and the reduction in metabolic age acceleration described above suggests that the molecular response to treatment may reflect this protection. Connecting a drug’s molecular effects to the diseases it prevents ordinarily requires large outcome trials. DynamiQ offers a more direct route. At the time of this analysis, the biorepository linked 65,847 plasma samples from 11,920 individuals to longitudinal clinical records. Because participants are sampled repeatedly over years, many contribute plasma collected before a first diagnosis, allowing features associated with *incident* disease to be identified across the full untargeted metabolome. Because the treated cohort was profiled on the same platform, its pharmacodynamic signature can be projected directly onto this disease landscape. We therefore asked whether the metabolite features that changed with treatment (Fig. 2a) are also associated with incident disease in DynamiQ, comparing plasma collected more than 90 days before diagnosis with plasma from age-and sex-matched healthy controls (Fig. 5a). For every incident outcome examined, treatment-responsive features overlapped with disease-associated features more than expected by chance (one-sided Fisher’s exact test, FDR < 10⁻⁹⁰ for each outcome; Supplementary Table 16). Because a feature can change with treatment in either direction relative to its association with disease, we focused on the overlapping features whose change opposed the disease association, increasing with treatment where higher levels marked lower odds of disease or decreasing where they marked higher odds. Such features were found for incident heart failure (788 features), liver fibrosis (739), type 2 diabetes (688), chronic kidney disease (536) and metabolic dysfunction-associated steatohepatitis (MASH; 489), as well as for Alzheimer’s disease (121) and Parkinson’s disease (92) (Fig. 5b; Supplementary Fig. S9). For the cardiometabolic, renal and hepatic outcomes, these features made up 28% (chronic kidney disease) to 56% (liver fibrosis) of the full overlap. The remaining overlapping features changed in the same direction as the disease association. Across all outcomes, including the neurodegenerative ones, the full overlap was 2.6-to 4.7-fold greater than expected by chance (Supplementary Table 16a). The treatment-responsive metabolome therefore overlaps extensively with the metabolic state that precedes cardiometabolic, renal and hepatic disease, and a substantial set of these features shifts away from the disease-associated state with treatment, but the overlap extends across outcomes rather than being confined to the diseases that incretin agonists are known to prevent, which led us to ask whether a second molecular layer resolves the disease associations more specifically.

**Figure 5.**
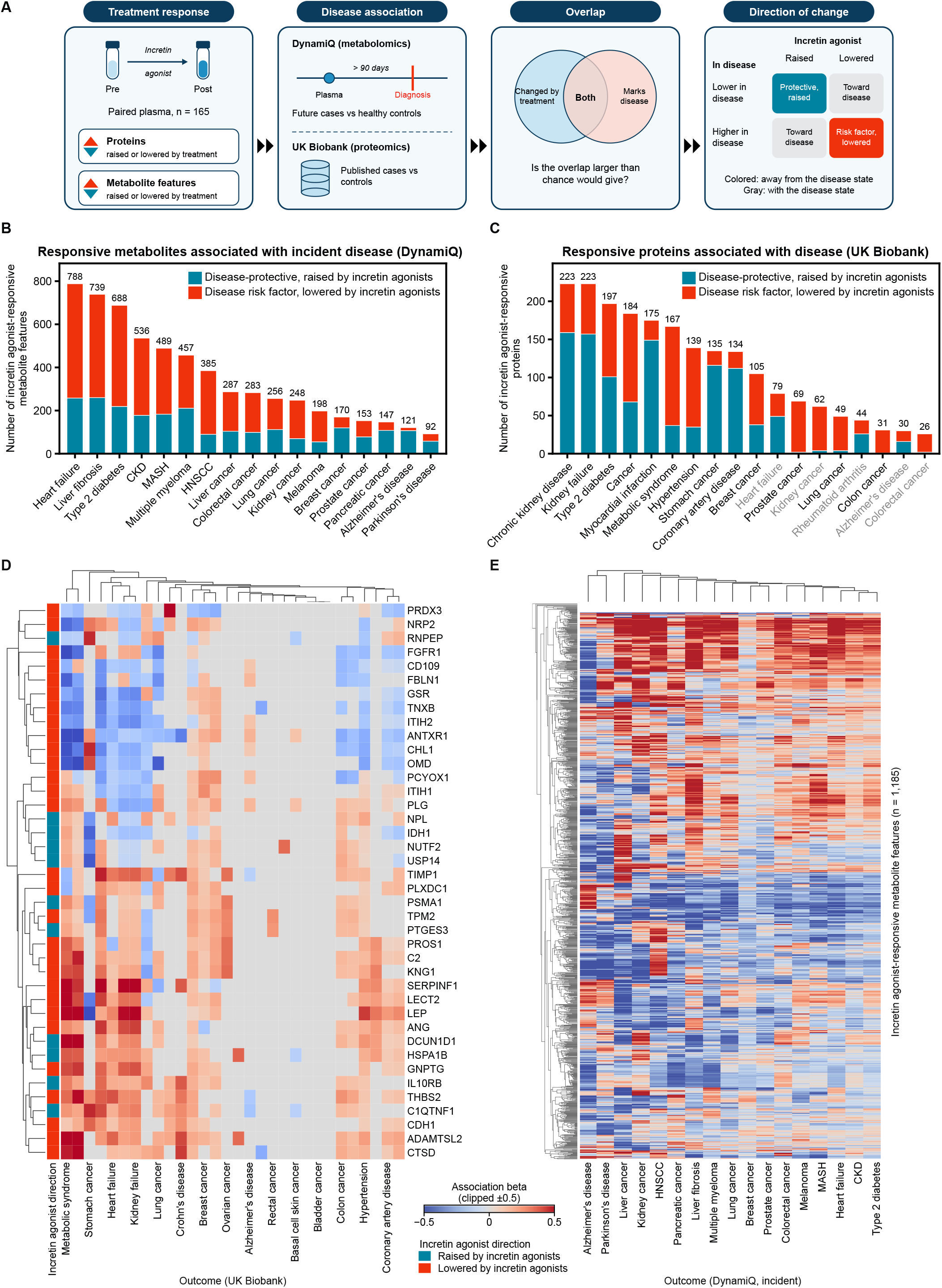
Incretin agonist-responsive features overlap with features of cardiometabolic, renal and cancer risk. a, Analysis design. Features that changed with treatment in the paired cohort (proteins and metabolite features) were intersected with features associated with disease, from pre-diagnostic plasma in DynamiQ (metabolomics) and from published UK Biobank case-control statistics (proteomics); the overlap was tested against all measured features, and overlapping features were classified by whether their change with treatment opposed (colored) or followed (gray) the direction of their association with disease. b, Number of incretin agonist-responsive metabolite features associated with incident disease outcomes in DynamiQ whose change with treatment opposed the disease association, divided into disease-protective features raised by treatment (teal) and disease risk factors lowered by treatment (red); fold enrichment of the full overlap, numbers of cases and controls, and features changing in the same direction as the disease association are given in Supplementary Table 16a. c, As in b, for treatment-responsive proteins against published UK Biobank case-control proteomic summary statistics (Supplementary Table 16b); outcomes whose overlap did not exceed chance (FDR ≥ 0.1) are labeled in gray. d, Hierarchically clustered heatmap of association effect sizes between 40 treatment-responsive proteins (rows) and disease outcomes (columns) in UK Biobank; the bar at left shows whether each protein was raised (teal) or lowered (red) by incretin agonists; dendrograms show clustering by Euclidean distance with average linkage; color scale indicates the association estimate, clipped at ±0.5, and gray cells have no estimate; outcome labels are shown for a subset of columns. e, Hierarchically clustered heatmap of association effect sizes between responsive metabolite features (rows; n = 1,185) and incident disease outcomes in DynamiQ (columns); color scale indicates the association estimate, clipped at ±0.5.

To test whether the disease mapping holds in an independent population and a second molecular layer, we repeated the analysis for the 1,080 treatment-responsive proteins against published UK Biobank proteomic case-control summary statistics^39^, and here the overlap was more selective. Treatment-responsive proteins whose change opposed the disease association were found for kidney failure and chronic kidney disease (223 proteins each), type 2 diabetes (197), myocardial infarction (175), metabolic syndrome (167), hypertension (139) and coronary artery disease (134) (Fig. 5c; Supplementary Fig. S10), and the full overlap with each of these outcomes exceeded chance (1.20-to 1.54-fold enrichment, FDR ≤ 2.7 × 10⁻⁵; Supplementary Table 16). By contrast, the overlap with proteins associated with heart failure (FDR = 0.14), Alzheimer’s disease (FDR = 0.98), cerebral infarction (FDR = 0.98) and rheumatoid arthritis (FDR = 0.32) did not exceed chance. Treatment-responsive proteins also overlapped with proteins associated with clinical laboratory measures of liver function, kidney function, glycemia and blood lipids in UK Biobank (Supplementary Fig. S11). In the proteome, therefore, the response to incretin agonists maps preferentially onto atherosclerotic cardiovascular, renal and metabolic disease, the conditions in which these drugs have shown benefit in outcome trials, and not onto neurodegenerative or autoimmune disease, so that a population biobank measured on a different platform recovers much of the established clinical profile of incretin agonists from molecular data alone.

Beyond the established indications, both molecular layers also overlapped with features associated with cancer. In the metabolome, treatment-responsive features whose change opposed the disease association were found for multiple myeloma (457 features), head and neck squamous cell carcinoma (385), liver cancer (287), colorectal cancer (283) and pancreatic cancer (147), and the overlap exceeded chance for every incident cancer examined (Fig. 5b; Supplementary Table 16). In the proteome, the overlap exceeded chance for cancer overall (184 proteins opposing the disease association; FDR = 7.0 × 10⁻¹⁷) and for stomach, prostate, colon, lung and breast cancer, but not for kidney, pancreatic or colorectal cancer (Fig. 5c; Supplementary Table 16). Multiple myeloma, hepatocellular, colorectal and pancreatic cancer are among the obesity-associated cancers whose incidence was lower in people with type 2 diabetes treated with GLP-1RAs than in those treated with insulin^40^. Obesity-associated cancers share metabolic and inflammatory drivers with the cardiometabolic diseases described above, and the overlap of the treatment response with cancer-associated features in both layers suggests that pharmacodynamic mapping against real-world longitudinal data can help prioritize candidate indications before outcome trials are designed.

The disease associations of treatment-responsive features were organized by disease biology. When the effect sizes linking treatment-responsive proteins to UK Biobank outcomes were clustered, proteins lowered by treatment formed a block positively associated with metabolic syndrome, heart failure and kidney failure, consistent with a shared set of treatment-responsive proteins associated with cardiometabolic and renal risk (Fig. 5d). The corresponding metabolite associations in DynamiQ showed the same organization: heart failure, chronic kidney disease, type 2 diabetes and MASH clustered together, while Alzheimer’s and Parkinson’s disease formed a separate group at the opposite end of the clustering (Fig. 5e). The treatment-responsive metabolome therefore carries distinct signatures for metabolic and neurodegenerative disease rather than a single generic disease signal. This resolution makes it possible to ask which parts of a drug’s response bear on which diseases.

Together, these results place the molecular response to incretin agonists within the biology of the cardiometabolic, renal and hepatic diseases these drugs prevent, most selectively in the proteome, and they extend it to emerging indications such as obesity-associated cancers. Because the treated cohort was measured on the same platform as a longitudinal biorepository that samples people before they are diagnosed, these observations also suggest a general approach for relating a pharmacological intervention to real-world disease risk, in which a drug’s pharmacodynamic signature is compared with the features that mark disease. The same logic can be applied in reverse. If the features that change with treatment are those that mark disease risk, then the state of these features before treatment describes the biology a drug must act on, and it may shape how much a patient benefits.

### Baseline inflammation panel profile is associated with subsequent weight loss

Weight loss on incretin agonists varied widely in this cohort, and 31 participants lost no weight despite treatment (Fig. 1a,b). Because treatment-responsive features correspond to indicators of real-world disease risk, we reasoned that differences in these features before treatment might distinguish participants who respond from those who do not. We compared baseline plasma between participants who went on to lose more than 5% of body weight and those who did not, across all three molecular layers. Neither the baseline metabolome (34,615 features) nor the baseline proteome (5,708 proteins) differed between these groups. By contrast, 38 of 248 inflammation panel proteins did (n = 147 participants with baseline measurements and weight follow-up), and every one of the 38 was lower at baseline in participants who went on to lose weight (Fig. 6a). Of the three molecular layers, only the inflammatory state of a patient before treatment carried information about how much weight they would subsequently lose. The baseline metabolome, which carries the strongest signatures of cardiometabolic disease risk (Fig. 5b), did not separate the groups. This suggests that the capacity to respond depends less on the degree of metabolic dysfunction at baseline than on the immune state that accompanies it.

**Figure 6.**
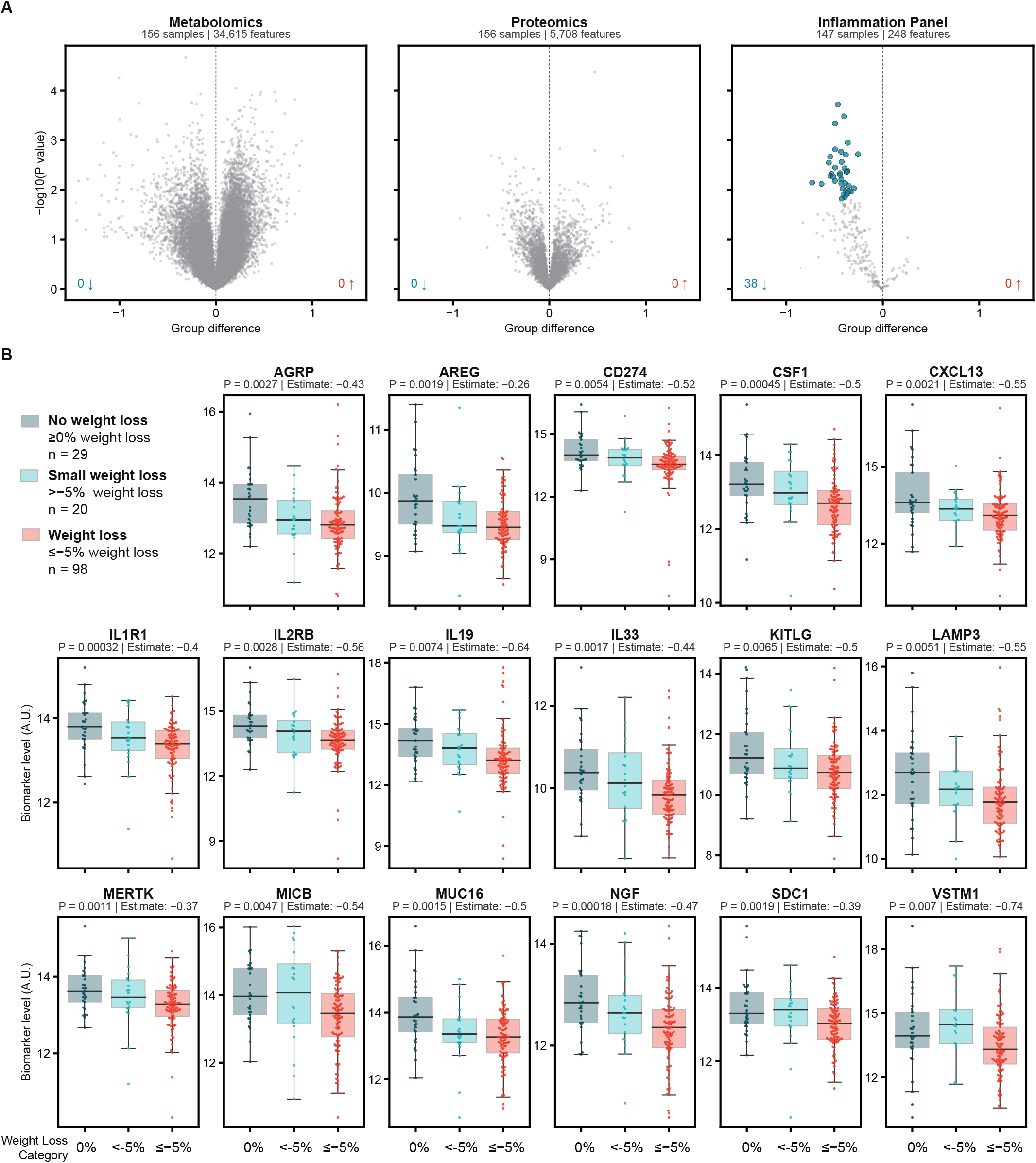
Baseline inflammation panel profile is associated with subsequent weight loss. a, Volcano plot of the baseline unpaired comparison between participants losing more than 5% of body weight and the remainder (n = 147 participants); all 38 significant analytes have negative estimates. b, Baseline levels of 17 of the 38 inflammation panel analytes that differed at baseline (a) across weight-loss response groups; group sizes are given in participants (no weight loss, n = 29; 0–5% loss, n = 20; ≥5% loss, n = 98); P values from one-way ANOVA; estimates from linear regression adjusted for age and sex.

This raises the question of what kind of immune state is associated with a weaker response to treatment. The analytes that distinguished the response groups did not resemble an acute inflammatory response, and instead described chronic tissue stress and ongoing repair (Fig. 6b). They included the alarmin IL-33 (P = 1.7 × 10⁻³), which is released by stressed stromal and epithelial cells^41^, and amphiregulin (AREG; P = 1.9 × 10⁻³), a repair factor that acts downstream of IL-33 signaling^42^. They also included the IL-1 family receptor IL1R1 (P = 3.2 × 10⁻⁴) and nerve growth factor (NGF; P = 1.8 × 10⁻⁴). Macrophage growth and activation was represented by CSF1 (P = 4.5 × 10⁻⁴), KITLG (P = 6.5 × 10⁻³), which treatment also lowered (Fig. 2a), and the efferocytosis receptor MERTK (P = 1.1 × 10⁻³). Ligands that stressed cells shed or display to the immune system were represented by MICB (P = 4.7 × 10⁻³), PD-L1 (CD274; P = 5.4 × 10⁻³) and syndecan-1 (SDC1; P = 1.9 × 10⁻³), together with the lymphoid chemokine CXCL13 (P = 2.1 × 10⁻³). Agouti-related protein (AGRP; P = 2.7 × 10⁻³) was also among them. AgRP is the orexigenic neuropeptide whose signaling incretin agonists suppress in the hypothalamus^43^, and it is a component of the proteomic aging clock described above^37^. Across the three weight-loss categories, levels of most of these analytes were highest in participants who lost no weight and lowest in those who lost the most (Fig. 6b). The association was most evident in participants over fifty years of age and in those who were not obese at baseline (Supplementary Fig. S12). Together, these analytes describe a state of chronic tissue stress, macrophage activation and repair, of the kind that characterizes inflamed adipose tissue in obesity, and a higher level of this state before treatment was associated with a smaller weight-loss response. The inflammation panel thus emerges as the layer most closely tied to individual variation in response. It was the only layer that distinguished participants before treatment, and, like the proteome, its response to treatment largely did not depend on the weight participants lost (Fig. 3a). A patient’s immune state therefore appears both to shape the response to incretin agonists and to be reshaped by it.

Taken across all three molecular layers, the response to incretin agonists in this real-world cohort resolves into two components that relate differently to weight loss and to disease. The first is a triacylglycerol-dominated lipid signature in the metabolome that scales with each participant’s weight loss (Fig. 3a,b), consistent with the mobilization of stored fat. The second is a broader response that does not depend on weight loss, spanning lower circulating mitochondrial oxidative proteins, higher levels of mTOR-axis regulators and a shift of the inflammation panel toward a less inflamed state (Fig. 2), and it is this broader response that accompanies the reduction of nearly three years in metabolic age acceleration (Fig. 4). The features that change with treatment overlap with features associated with incident cardiometabolic, renal, hepatic and obesity-associated cancers in DynamiQ, and in UK Biobank they overlap most selectively with atherosclerotic cardiovascular, renal and metabolic disease (Fig. 5), the conditions in which these drugs have shown clinical benefit. Immune state runs through these observations, because inflammation-panel mediators of tissue remodeling and innate immunity were associated with the reduction in metabolic age (Fig. 4h), and the inflammatory state present before treatment was associated with how much weight participants went on to lose (Fig. 6). Together, these results describe a response to incretin agonists that extends well beyond weight loss, overlaps with the molecular features of the diseases these drugs are known to reduce, and is shaped by the inflammatory state in which treatment begins.

Across the three molecular layers, the extent to which the response depended on weight loss differed sharply. Weight loss correlated with 596 metabolite features, dominated by triacylglycerols, but with only 1 protein and 14 inflammation panel analytes (Fig. 3a,b). The broader response, which did not depend on weight loss, accompanied a 2.96-year reduction in metabolic age acceleration that was itself unrelated to weight loss (Fig. 4d,f). Treatment-responsive features overlapped with features associated with all 17 incident outcomes in DynamiQ (2.6-to 4.7-fold enrichment) and, in UK Biobank, selectively with renal, metabolic and atherosclerotic cardiovascular disease (Fig. 5). The inflammation panel connected these findings: changes in five of its analytes were associated with the change in metabolic age (Fig. 4h), and 38 of its analytes were lower at baseline in participants who went on to lose more than 5% of body weight (Fig. 6a). Together, these findings indicate that much of the molecular response to incretin agonists does not depend on weight loss and that inflammatory state is associated with both the response to treatment and its magnitude.

## Discussion

Incretin agonists produce substantial weight loss and reduce the risk of cardiovascular, renal and metabolic disease, yet the extent to which their benefits depend on weight loss itself remains unclear. It is also unclear why individuals receiving the same treatment differ so markedly in their response, and whether these drugs act on the biology of aging. To address these questions, we profiled paired plasma samples collected before and after treatment in a real-world cohort across the proteome and the metabolome, and related the changes to weight loss, metabolic age and disease risk. Treatment was associated with broad remodeling of the plasma proteome and metabolome, and most of this remodeling did not depend on the weight participants lost. That included a reduction in circulating mitochondrial oxidative proteins, a shift toward a less inflamed state and a decrease of nearly three years in metabolic age acceleration. The features that changed with treatment overlapped with features associated with cardiometabolic, renal and hepatic disease, consistent with the established clinical benefits of these drugs, and before treatment, higher systemic inflammation was associated with less subsequent weight loss. Together, these findings suggest that incretin agonist treatment is associated with a reduction in the tissue stress that accompanies obesity, a reduction that does not depend on weight loss, and that the same stress, when pronounced before treatment, may limit how much weight a patient loses.

Most of the molecular response to incretin agonists did not depend on weight loss, but the degree of this dependence differed between molecular layers. The metabolome carried a measurable signature of weight loss after incretin treatment, dominated by triacylglycerols that scaled with the weight lost, whereas the proteomic and inflammatory changes were largely unrelated to it. A previous proteomic study of semaglutide found that a subset of protein changes persisted after statistical adjustment for weight loss^17^. By measuring three molecular layers in the same participants, we show that the relationship to weight loss depends on the layer: the metabolome correlates with weight loss, whereas the proteome and inflammation panel largely do not. Because this conclusion rests on the absence of a correlation with weight loss, it indicates that these changes did not depend on the extent of weight loss in this cohort, not that weight loss plays no part in them. The most prominent change not dependent on weight loss was the decrease in circulating mitochondrial oxidative proteins and cardiolipins. In obesity and insulin resistance, hepatic mitochondria operate at elevated capacity in response to excess lipid supply^28^, and the rodent evidence described above indicates that incretin-based treatment can lower this load in ways that food restriction does not^29^. In our cohort, the mitochondrial decrease occurred without any increase in the cytokines and receptors that signal tissue injury, and FGF21, a marker of mitochondrial stress, fell. These observations point to relief of lipid-driven mitochondrial overload, and with it a reduction in tissue stress.

Obesity is a chronic inflammatory state^44^, and weight loss is generally thought to resolve that inflammation. Our data suggests that the relationship runs in both directions. After treatment, the inflammation panel shifted toward a less inflamed state in a way that largely did not depend on the weight participants lost, indicating that the lower systemic inflammation during treatment is not simply a consequence of weight loss. Before treatment, by contrast, a higher inflammatory state was associated with less weight loss. The analytes that distinguished participants who went on to lose weight did not resemble an acute inflammatory response. Instead, they described chronic tissue stress, macrophage activation and repair, including IL-33, amphiregulin, CSF1, MERTK and the stress-induced ligand MICB, a state characteristic of inflamed adipose tissue in obesity. These findings are consistent with a model in which pre-existing inflammation limits the weight-loss response, potentially by blunting the central and peripheral signaling through which these drugs act, much as the hypothalamic inflammation and gliosis that accompany obesity are thought to impair central regulation of energy balance^45^. In a dietary weight-loss intervention, a comparably sized inflammatory and cardiovascular protein panel identified only a single baseline predictor^46^, suggesting that inflammatory state is particularly relevant to the response to incretin agonists. If obesity and inflammation reinforce each other in this way, relying on weight loss to resolve inflammation may be insufficient for patients whose inflammatory burden blunts their response. This raises the question of whether obesity and its accompanying inflammation should be treated together, for example by combining incretin agonists with therapies that target chronic tissue inflammation.

The reductions in mitochondrial and inflammatory stress that did not depend on weight loss bear directly on aging. Aging is marked by the accumulation of chronic cardiometabolic, renal and hepatic disease and the tissue stress that accompanies it, and it is increasingly regarded as modifiable^47^. Like the mitochondrial and inflammatory changes, the reduction in metabolic age acceleration did not depend on weight loss. It was greatest in participants who began with the greatest age acceleration, most of whom had diabetes, indicating that the changes associated with treatment are most pronounced where metabolic burden is high. These observations suggest that baseline metabolic state shapes the magnitude of the response to incretin agonists, such that patients starting from different metabolic set points may derive different degrees of benefit from treatment. Because participants with diabetes outnumbered those without by roughly two to one, larger cohorts of people without diabetes will determine whether their response differs in kind or mainly in magnitude, and how much improved glycemia contributes. Participants whose growth-signaling and immune pathways rose most, and whose tissue-remodeling mediators such as OSM, MMP9, MMP8 and PTX3 increased most, showed the largest decreases in age acceleration. Metabolic and inflammatory burden therefore appear to play different roles. Greater metabolic burden was associated with a larger reduction in metabolic age, whereas greater inflammatory burden was associated with less weight loss. The reduction in metabolic age is consistent with multi-omic changes in aged mice^12^, with slower epigenetic aging in semaglutide-treated people with HIV-associated lipohypertrophy^14^, and with the decrease in three components of a UK Biobank proteomic aging clock in our cohort^37^. The link to age-related disease was also visible in the treatment-responsive features themselves. Because the cohort was profiled on the same platform as a biorepository that samples individuals before diagnosis, we could relate these features to incident and prevalent disease. They aligned with the established clinical benefits of these drugs, and also with obesity-associated cancers whose incidence is lower in patients treated with GLP-1RAs than in those treated with insulin^40^. Although these associations come from external populations, relating pharmacodynamic changes to disease risk in this way may help generate hypotheses about additional indications.

Baseline inflammatory disposition is one source of the marked heterogeneity in response to these drugs, and our data suggest it is not the only one. Hierarchical clustering of individual responses in the 155 participants with complete data revealed distinct molecular subgroups that demographic categories alone do not capture (Supplementary Fig. S13). Efforts to predict who will respond have so far focused largely on germline genetic variation. The largest genome-wide association study of weight loss with these drugs, in 27,885 users, identified a single significant locus in GLP1R, associated with 0.76 kg of additional weight loss per allele, and genetic factors added little to the variance explained by demographic and clinical factors alone^48^. Studies of glycemic response have likewise found only modest-effect variants^49^. In contrast, we identified 38 baseline inflammatory mediators associated with subsequent weight loss in 147 participants. Differences in design preclude a direct comparison, but the contrast suggests that circulating inflammatory mediators capture determinants of response that germline variation does not. Validating the baseline inflammatory signature in independent cohorts, characterizing these response subgroups and comparing genomic and circulating predictors within a single cohort are important directions for future work. The approach taken here may be useful wherever the response to a therapy varies between patients. Profiling the same individuals before and after treatment, stratifying them by clinical response and relating the treatment-associated features to disease risk in a biorepository profiled on the same platform yielded pharmacodynamic markers separable from weight loss, a baseline inflammatory state associated with a weaker response, and hypotheses about additional indications, each of which corresponds to a recurring need in drug development. Pathways identified here, such as chronic inflammatory mediators that marked weaker responders, also suggest candidates for combination with incretin agonists, although each of these leads requires validation in independent cohorts.

In summary, this study resolves the response to incretin agonists into a lipid profile that scales with weight loss and a broader profile spanning mitochondrial metabolism, inflammation and growth signaling, which together accompany a reduction in metabolic age. Matched untreated cohorts would strengthen the conclusion that the reduction in metabolic age reflects treatment. Because the cohort lacked an untreated comparison group, the changes that did not depend on weight loss are best described as associated with treatment, and time-matched comparisons with repeated samples from untreated individuals in the same biorepository offer a direct way to establish which of these changes are specific to treatment. Measurements of body composition would further resolve how lean and fat mass contribute to these two profiles, and prospective studies would extend the disease associations, drawn here from prevalent and incident disease associations in other individuals, to clinical outcomes. Beyond these next steps, the findings carry implications that reach past weight management. They suggest that the biology of aging can be modulated by relieving the metabolic and inflammatory stress of obesity, in ways that do not depend on weight loss alone. They indicate that a patient’s inflammatory profile before treatment could help identify those likely to benefit from incretin agonists alone and those who may need combination therapy. And they show that real-world biorepositories linked to longitudinal multi-omic data can relate the pharmacodynamics of a therapy to the diseases it may prevent, offering a route to understanding how established and emerging treatments act in the patients who receive them.

## Summary, Impact and Limitations

Herein we sought to determine how much of the molecular response to incretin agonists depends on weight loss. The analysis resolves that response into two profiles: a lipid profile dominated by triacylglycerols that scaled with the weight participants lost, and a broader profile spanning mitochondrial metabolism, inflammation and growth signaling that did not depend on weight loss and was accompanied by a reduction of nearly three years in metabolic age acceleration. Treatment-associated features overlapped with features associated with cardiometabolic, renal and hepatic disease, and a higher inflammatory state before treatment was associated with less subsequent weight loss. These findings suggest that lower metabolic and inflammatory stress during treatment may be accompanied by slower biological aging, and that a patient’s inflammatory profile could help distinguish those likely to benefit from incretin agonists alone from those who may need combination therapy, with INHBE^32–34^ and the chronic inflammatory mediators that marked weaker responders as candidate partners. More broadly, profiling the same individuals before and after treatment and relating their response to disease risk in a biorepository profiled on the same platform offers an approach that may be applied to any therapy whose effects vary between patients.

These conclusions carry the caveats that accompany any analysis of real-world treatment. Paired samples were collected in routine care, so changes between them may also reflect elapsed time and changes in concurrent care, and we therefore describe the changes that did not depend on weight loss as associated with treatment rather than as direct effects of the drug; repeated samples from untreated individuals in the same biorepository will allow time-matched comparisons that identify which of these changes are specific to treatment. The molecular response was concentrated in participants with type 2 diabetes, who outnumbered those without diabetes by roughly two to one (Supplementary Table 2), so improved glycemia may contribute to it, and the response of people without diabetes remains to be defined in larger cohorts. The disease overlaps relate treatment-associated features to risk measured in other individuals and are best read as hypothesis-generating. Because incretin agonists are prescribed largely to people with type 2 diabetes, the treated cohort shares a diabetic state with many of the disease cases in DynamiQ but not with the healthy controls, who were defined in part by the absence of diabetes; in an observational design this confounding is inherent and cannot be removed by matching, and it likely contributes to the breadth of the metabolomic overlap across outcomes, so the more selective overlap in the UK Biobank proteome carries more weight for disease specificity. Finally, the baseline inflammatory signature, identified here in a single cohort, requires validation in independent cohorts before it can guide the selection of patients for treatment. Answering these questions would move treatment from regarding obesity as a single condition toward matching each patient with the therapy, or combination of therapies, best suited to the biological state from which they begin.

## Methods

### Study cohort

The study cohort was defined by querying the Sapient DynamiQ biorepository of human plasma samples linked with multi-omics and real-world data for participants prescribed a GLP-1-based therapy (semaglutide, liraglutide, dulaglutide or tirzepatide) for at least 90 days. Tirzepatide, a dual GIP/GLP-1 receptor co-agonist, accounted for just over a quarter of the cohort (47 of 165 participants), either alone or in combination with another agent; no participants had recorded exposure to exenatide or lixisenatide. Of the 165 participants, 41 (25%) had a treatment record indicating exposure to more than one GLP-1-based agent during the study period (for example, a switch or combination between semaglutide and tirzepatide), rather than continuous single-agent monotherapy throughout. For each participant, the plasma sample closest to but prior to the first recorded prescription date was selected as the pre-treatment timepoint, and the sample closest to but prior to the last recorded prescription date was selected as the post-treatment timepoint.

Percent weight loss was calculated from electronic health record weight measurements closest to each sample date and used to classify participants into three response groups: no weight loss (weight change ≥ 0%), small weight loss (loss of less than 5%) and weight loss (loss of 5% or more, that is, a change of −5% or below). A subset of baseline predictor analyses (Fig. 6) used a strict binary threshold (weight loss >5% versus ≤5%) rather than this three-category grouping. The specific grouping used is noted for each analysis below, and the two definitions yield slightly different group sizes due to differences in the weight measurement timepoint used for each. Of the 165 treated participants, 155 had inflammation panel measurements at both time points, and 156 had sufficient weight data to be assigned to a response group (no weight loss, n = 31; 0–5% loss, n = 22; ≥5% loss, n = 103); analyses involving both weight and the inflammation panel used the 147 participants with both (n = 29, 20 and 98, respectively). Baseline obesity was defined as body mass index ≥ 30 kg m⁻². Inclusion required at least 90 days between paired samples, at least 90 days of total treatment, and weight measurements within 9 months before and 3 months after treatment initiation. No minimum or maximum age restriction was applied for cohort eligibility. Participant age at baseline ranged from 15.7 to 82.3 years, including one adolescent female participant (age 15.7 years, with a diabetes diagnosis) whose participation was covered under IRB-approved consent procedures, including guardian consent and participant assent (see Ethics declaration). These criteria yielded a final cohort of 165 participants (330 paired samples) from an initial population of 11,920 participants.

### Sample collection and multi-omic profiling

Plasma samples (K2-EDTA) were collected, stored at –80°C and processed blinded to clinical metadata. Proteomics was performed by magnetic nanoparticle-based protein enrichment, on-bead tryptic digestion, and data-independent acquisition mass spectrometry (Bruker timsTOF HT), with peptides identified against a canonical human proteome using directDIA (Spectronaut) at 1% false discovery rate. Non-targeted metabolomics was performed by rapid liquid chromatography–mass spectrometry (rLC-MS) in positive and negative ionization modes (Bruker timsTOF Pro 2), with feature extraction, drift correction and annotation against an internal spectral library. Cytokine profiling used a targeted, sequencing-based immunoassay panel (NULISAseq, Alamar Biosciences) on the ARGO-HT platform. Full reagent, instrument and quality control details are provided in Supplementary Methods.

### Statistical analysis

All analyses were performed using an internal analysis package built on Python and scikit-learn^50^. Metabolomics data were filtered for isotopologues, prevalence and intensity. All omics layers were prevalence-filtered, imputed, log-transformed and standardized prior to analysis. Pharmacodynamic biomarkers were identified by two-sided paired t-tests comparing pre-and post-treatment values, performed across the full cohort and within strata defined by sex, age (above or below 50 years), diabetes status and baseline obesity. Association between biomarker fold change and percent weight loss was assessed by Spearman correlation and by ordinary least squares regression adjusted for sex, age, diabetes status and baseline body mass index, in the full cohort and within the same strata. Baseline associations with subsequent weight loss were assessed in two ways: by unpaired t-test between participants above and below a 5% weight loss threshold, and by regression of baseline level against a continuous weight loss outcome; the test used is stated for each result. Multiple testing was corrected using the Benjamini-Hochberg procedure at a false discovery rate of 0.1, applied separately to annotated and unannotated metabolomic features. Between-group differences in the within-person change in age acceleration were tested by ordinary least squares regression of the change on the group indicator, with and without baseline age acceleration as a covariate. Permutation testing of the sex-stratified baseline proteomic analysis used 1,000 shuffles of the weight loss outcome with the full analysis pipeline re-run on each shuffle; stability was assessed by leave-one-participant-out resampling (Supplementary Table 17).

## Age acceleration analysis

A metabolic aging clock was retrained on the present dataset following the methodology described previously for this platform^36^, using Lasso regression with hyperparameters tuned by 5-fold cross-validation (Optuna)^51^ on healthy control participants from the same biorepository, split 70/30 at the participant level into training and held-out sets. Model predictions were then detrended against chronological age using a linear regression fitted on the training set, with detrended prediction defined as (prediction - intercept) / slope; the fitted intercept was 14.66 and the slope 0.69. Because the slope is less than one, this transformation rescales age acceleration by a factor of 1/0.69 = 1.45, inflating its standard deviation by approximately 45% and its variance by approximately 110% relative to the raw predictions, which should be borne in mind when interpreting the range of age acceleration shown in Fig. 4. Age acceleration was defined for each sample as the difference between detrended model-predicted biological age and chronological age. Pre-and post-treatment age acceleration values were paired per participant and the within-person change (post minus pre) assessed by two-sided paired t-test across the full cohort and within strata, summarized as a forest plot of the mean within-person change and its 95% confidence interval (Fig. 4e). Two participants fell outside the chronological age range of the training set; a sensitivity analysis excluding them is reported in Supplementary Table 18. Sensitivity of the change to elapsed time was assessed by Spearman correlation and linear regression of the within-person change on the interval between paired samples, on exposure duration at the post-treatment sample, and on the calendar date of the baseline sample.

To relate the clock result to the measured proteome, a per-participant response score was computed for each of twelve protein gene sets as the mean z-scored per-participant log2 fold change of its member proteins, and correlated with the within-person change in age acceleration by Spearman correlation. Because the change is itself predicted by its baseline value, each correlation was repeated as a partial Spearman correlation and as an ordinary least squares regression with baseline age acceleration as a covariate. To separate pathway-specific from global effects, each participant’s overall response magnitude, defined as the mean absolute log2 fold change across responsive proteins, was added as a further covariate (Supplementary Table 19).

## Disease association and network analyses

Overlap between pharmacodynamic metabolite biomarkers and disease-relevant features was assessed in DynamiQ for both prevalent and incident disease, and overlap for protein biomarkers was assessed against published UK Biobank case-control proteomic summary statistics. DynamiQ associations were estimated by logistic regression per feature adjusted for age and sex, with cases and healthy controls matched by age and sex distribution. For each outcome, enrichment of treatment-responsive features among disease-associated features was tested by one-sided Fisher’s exact test against all tested features, with Benjamini-Hochberg correction across outcomes, and fold enrichment was calculated as the observed overlap divided by the overlap expected by chance. Overlapping features were further classified by whether their change with treatment was opposite to, or in the same direction as, their association with disease.

Correlation networks comparing pharmacodynamic biomarkers whose change did and did not depend on weight loss were constructed from pairwise Spearman correlations, retaining edges with absolute r > 0.7. Gene set enrichment of the protein set not dependent on weight loss was tested by one-sided Fisher’s exact test against all measured proteins across 104 gene sets, with Benjamini-Hochberg correction applied across gene sets (Supplementary Table 20).

## Ethics declaration

This study was conducted using de-identified plasma samples and linked clinical data from the DynamiQ biorepository. Demographic information such as age, gender, race, and ethnicity was collected at enrollment using standardized questionnaires and linked to each sample in the database. Clinical information was obtained from electronic health records and integrated with molecular measures generated in each sample using Sapient’s rLC-MS platform to enable deep phenotyping approaches and stratification of individuals into subgroups. Data handling and storage procedures were designed to protect the confidentiality and privacy of participant data and comply with all relevant regulatory rules and guidelines.

## Data availability

Individual-level metabolomics, proteomics and cytokine data generated in this study, together with associated de-identified clinical metadata, are held under data use agreements that restrict public deposition of individual-level data. Processed summary statistics supporting the findings of this study are provided in the Supplementary Information. Requests for access to de-identified individual-level data may be directed to the corresponding author. UK Biobank proteomic summary statistics used for disease association comparison are available from the published UK Biobank proteomic disease-association study^39^.

## Code availability

Code used for the analyses described in this manuscript is available from the corresponding author upon reasonable request.

## Acknowledgements

This work was funded by Sapient Bioanalytics.

## Author contributions

T.L. and M.J. conceived and designed the study. K.G., K.D. and S.P. performed the experiments. T.L. and A.L. analyzed the data. T.L., M.J. and J.L.J. wrote the draft. All authors reviewed and edited the manuscript. J.U., J.W. and M.J. supervised the study.

## Supplementary Figure Legends

**Supplementary Figure S1**

Demographic characteristics of the treated cohort. a, Distribution of age at baseline (n = 165 participants). b–e, Proportions of participants by gender (b), race (c), ethnicity (d) and age above 50 years (e).

**Supplementary Figure S2**

Mitochondrial proteins and injury-responsive inflammation-panel analytes after treatment. a, Log2 fold change (post minus pre) of the mitochondrial proteins named in the main text, grouped by respiratory chain complex (CI–CIV), ATP synthase, tricarboxylic acid cycle and β-oxidation (n = 165 participants; Supplementary Table 3); color scale clipped at ±0.5. b, Mean within-person change (post minus pre) in standardized level for 43 inflammation-panel analytes that respond to tissue injury, grouped as acute inflammatory cytokines, damage-sensing receptors, death ligands and their receptors, macrophage and clearance markers, and stress hormones (n = 155 participants with paired measurements; Supplementary Table 5); color scale clipped at ±0.5 standard deviations. In both panels, P values are from two-sided paired t-tests with Benjamini–Hochberg correction across all features measured on each platform, and asterisks denote FDR < 0.1.

**Supplementary Figure S3**

Calibration of the metabolic aging clock and distribution of the change in age acceleration. a, Clock predictions before detrending versus chronological age in the training set (190 samples from 129 participants); the solid line shows the linear fit used for detrending (slope 0.69, intercept 14.66) and the dashed line shows identity, with Pearson r, R², root mean squared error and mean absolute error (95% confidence intervals in brackets) shown above the panel. b, The clock applied to the treated cohort, formatted as in a (Pearson r = 0.47, R² = −1.12). c, Distribution of the within-person change in age acceleration (post minus pre) in participants with diabetes (teal, n = 113) and without diabetes (red, n = 52), shown as the proportion of participants in each group; the dashed line marks no change. d, e, Number of participants by change in age acceleration among those with diabetes (d) and without diabetes (e).

**Supplementary Figure S4**

Molecular correlates of the change in age acceleration. a, Spearman correlation between the fold change of each treatment-responsive protein and the within-person change in age acceleration (n = 165 participants); colored points denote FDR < 0.1 and the most significant proteins are labeled. b, Within-person change in age acceleration versus participant-level pathway scores for mTORC1 signaling, PI3K/AKT/mTOR signaling, innate immune response and complement, where each score is the mean z-scored change of the pathway’s treatment-responsive proteins. Blue points and lines show the unadjusted relationship, and red points and lines show the relationship with both axes residualized on baseline age acceleration; panel headers give the unadjusted and partial Spearman r with FDR.

**Supplementary Figure S5**

Gene-set enrichment among treatment-responsive features with the inflammation panel analytes included alongside the proteins. a, Enrichment of gene sets among features that changed between paired pre-and post-treatment samples in all participants, formatted as in Fig. 2b; bars show the number of members with negative (blue, left) or positive (red, right) estimates, and FDR from one-sided Fisher’s exact test is shown at right, with asterisks denoting FDR < 0.1. b, The same analysis within strata defined by sex, diabetes status, age above or below 50 years and baseline obesity; panel headers name the stratum, and strata with no enriched gene sets are omitted.

**Supplementary Figure S6**

Pre-and post-treatment differences within demographic and clinical strata. Paired t-test volcano plots for metabolomics, proteomics and the inflammation panel within strata of sex, diabetes status, age above or below 50 years and baseline obesity, formatted as in Fig. 2a. Panel headers name the stratum and give the number of samples (two per participant). Blue points, significantly decreased; red points, significantly increased (FDR < 0.1); gray points, not significant; counts of decreased and increased features are shown in each panel.

**Supplementary Figure S7**

Enrichment of metabolite classes and protein gene sets within strata. a, Metabolite compound classes enriched among features that changed with treatment within each stratum, formatted as in Fig. 2c. b, Protein gene sets enriched among proteins that changed with treatment within each stratum, formatted as in Fig. 2b. Panel headers name the stratum, and strata with no enriched classes or gene sets are omitted.

**Supplementary Figure S8**

Relationship between feature fold change and weight change within strata. Volcano plots of the regression of each feature’s fold change on percent weight change for metabolomics, proteomics and the inflammation panel within the strata shown in Supplementary Fig. S6, formatted as in Fig. 3a. Panel headers name the stratum and give the number of samples.

**Supplementary Figure S9**

Overlap of treatment-responsive metabolite features with disease associations in DynamiQ. Each panel shows the paired pre-and post-treatment volcano plot of the metabolome, with features that are also associated with the indicated outcome highlighted (teal), shown separately for prevalent and incident disease across 15 outcomes.

**Supplementary Figure S10**

Overlap of treatment-responsive proteins with disease associations in UK Biobank. Each panel shows the paired pre-and post-treatment volcano plot of the proteome, with proteins that are also associated with the indicated outcome in published UK Biobank proteomic case-control summary statistics highlighted (teal), across 30 outcomes.

**Supplementary Figure S11**

Overlap of treatment-responsive proteins with associations with clinical laboratory measures in UK Biobank. Each panel shows the paired pre-and post-treatment volcano plot of the proteome for alanine transaminase, aspartate transaminase, estimated glomerular filtration rate, gamma-glutamyltransferase, HbA1c, high-and low-density lipoprotein cholesterol and triglycerides. Proteins associated with the measure are colored by whether the direction of their change with treatment matches (teal) or opposes (red) the direction of their association with the measure.

**Supplementary Figure S12**

Baseline differences by subsequent weight loss within strata. Unpaired t-test volcano plots comparing baseline levels between participants who went on to lose more than 5% of body weight and the remainder, for metabolomics, proteomics and the inflammation panel within strata of sex, diabetes status, age above or below 50 years and baseline obesity, formatted as in Fig. 6a. Panel headers name the stratum and give the number of participants.

**Supplementary Figure S13**

Participant-level correlation of molecular response profiles. Hierarchically clustered matrix of Spearman correlations between the fold-change profiles of individual participants.

## Supplementary Table Legend

**Supplementary Tables 1–20.** Cohort characteristics and statistical results. Unless stated otherwise, pre/post comparisons are two-sided paired t-tests; the estimate is the within-person change (post minus pre); corrected P values are Benjamini–Hochberg FDR, significant at FDR < 0.1. 1, cohort characteristics. 2, inventory of all analyses with sample, feature and hit counts. 3–5, pre/post changes in proteins (3, significant only; 1,080 protein features collapsing to 1,061 genes), annotated metabolites (4, significant only) and all 248 inflammation panel analytes (5, with the injury groups used in Supplementary Fig. S2b). 6, mTOR-axis components. 7, activin/TGF-β, IGF, muscle and adipokine axes, including undetected members. 8, aging-clock fit in training and held-out sets. 9, change in age acceleration by stratum. 10, change versus sampling interval, exposure duration and baseline date. 11, hemolysis and storage-time checks. 12, pathway scores versus change, unadjusted and baseline-adjusted. 13, per-protein correlation with change. 14, inflammation panel analytes associated with change. 15, between-group contrasts. 16, overlap of treatment-responsive features with disease associations: metabolites against prevalent and incident disease in DynamiQ, with case and control numbers (16a), and proteins against UK Biobank case-control associations (16b), each with expected overlap, fold enrichment, one-sided Fisher’s exact P and FDR, and the overlap split by direction relative to the disease association. 17, permutation and stability testing of the sex-stratified baseline proteomic analysis (negative result). 18, stratified change excluding two participants outside the clock training age range. 19, global response-magnitude term across pathway models. 20, gene-set enrichment of proteins not dependent on weight loss (negative result).

