## Supplementary Figures S1-S13 for "Real-world longitudinal multi-omics separates weight-loss-associated responses to incretin agonists from a broader response not dependent on weight loss"

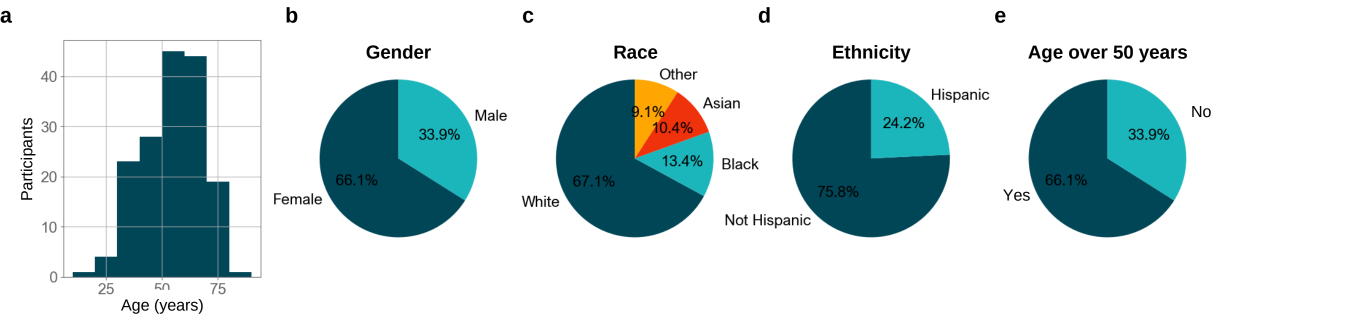
a
b
c
d
e
Participants
Age (years)
0
10
20
30
40
25
50
75
Gender
Male
Female
33.9%
66.1%
Race
Other
Asian
Black
White
9.1%
10.4%
13.4%
67.1%
Ethnicity
Hispanic
Not Hispanic
24.2%
75.8%
Age over 50 years
No
Yes
33.9%
66.1%

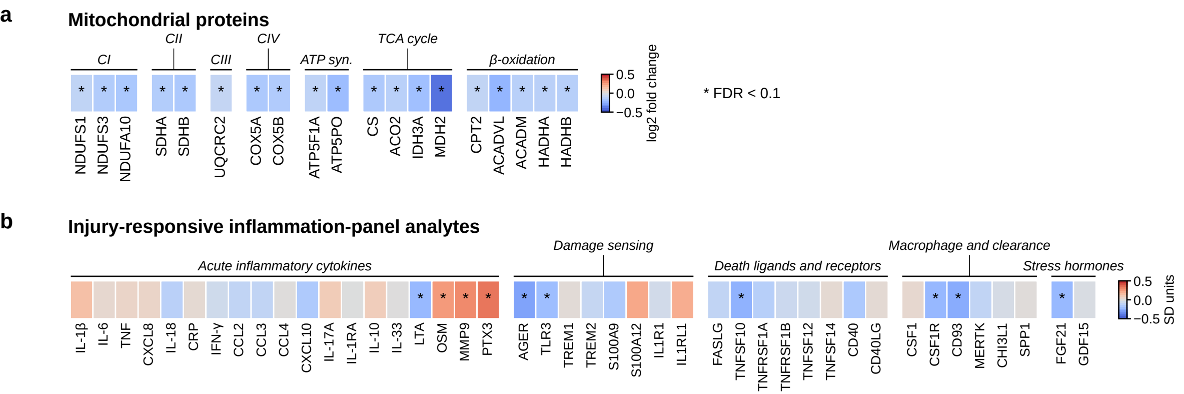
a
Mitochondrial proteins
CI
CII
CIII
CIV
ATP syn.
TCA cycle
β-oxidation
NDUFS1
NDUFS3
NDUFA10
SDHA
SDHB
UQCRC2
COX5A
COX5B
ATP5F1A
ATP5PO
CS
ACO2
IDH3A
MDH2
CPT2
ACADVL
ACADM
HADHA
HADHB
0.5
0.0
−0.5
log2 fold change
* FDR < 0.1
b
Injury-responsive inflammation-panel analytes
Acute inflammatory cytokines
Damage sensing
Death ligands and receptors
Macrophage and clearance
Stress hormones
IL-1β
IL-6
TNF
CXCL8
IL-18
CRP
IFN-γ
CCL2
CCL3
CCL4
CXCL10
IL-17A
IL-1RA
IL-10
IL-33
LTA
OSM
MMP9
PTX3
AGER
TLR3
TREM1
TREM2
S100A9
S100A12
IL1R1
IL1RL1
FASLG
TNFSF10
TNFRSF1A
TNFRSF1B
TNFRSF12
TNFSF14
CD40
CD40LG
CSF1
CSF1R
CD93
MERTK
CHI3L1
SPP1
FGF21
GDF15
0.5
0.0
−0.5
SD units

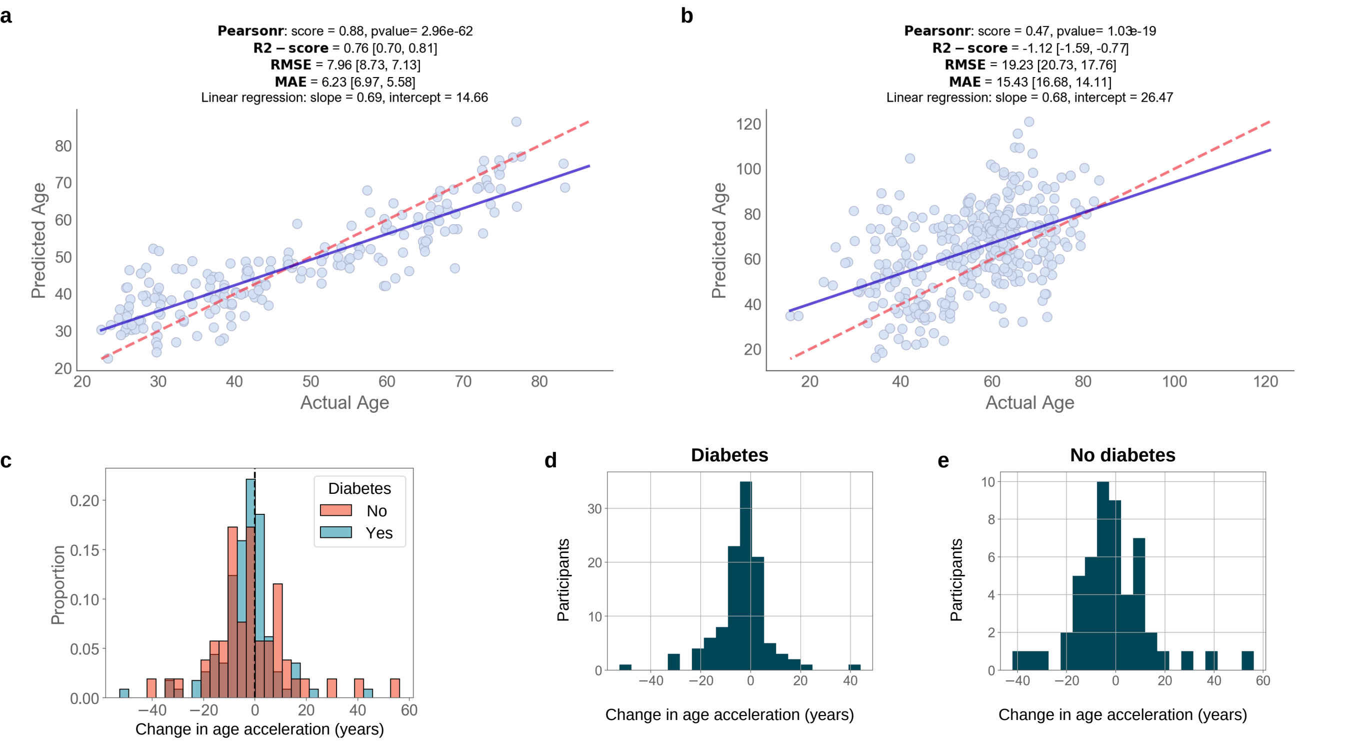
a
Pearsonr: score = 0.88, pvalue= 2.96e-62
R2 – score = 0.76 [0.70, 0.81]
RMSE = 7.96 [8.73, 7.13]
MAE = 6.23 [6.97, 5.58]
Linear regression: slope = 0.69, intercept = 14.66
Predicted Age
Actual Age
b
Pearsonr: score = 0.47, pvalue= 1.03e-19
R2 – score = -1.12 [-1.59, -0.77]
RMSE = 19.23 [20.73, 17.76]
MAE = 15.43 [16.68, 14.11]
Linear regression: slope = 0.68, intercept = 26.47
Predicted Age
Actual Age
c
Diabetes
No
Yes
Proportion
Change in age acceleration (years)
d
Diabetes
Participants
Change in age acceleration (years)
e
No diabetes
Participants
Change in age acceleration (years)

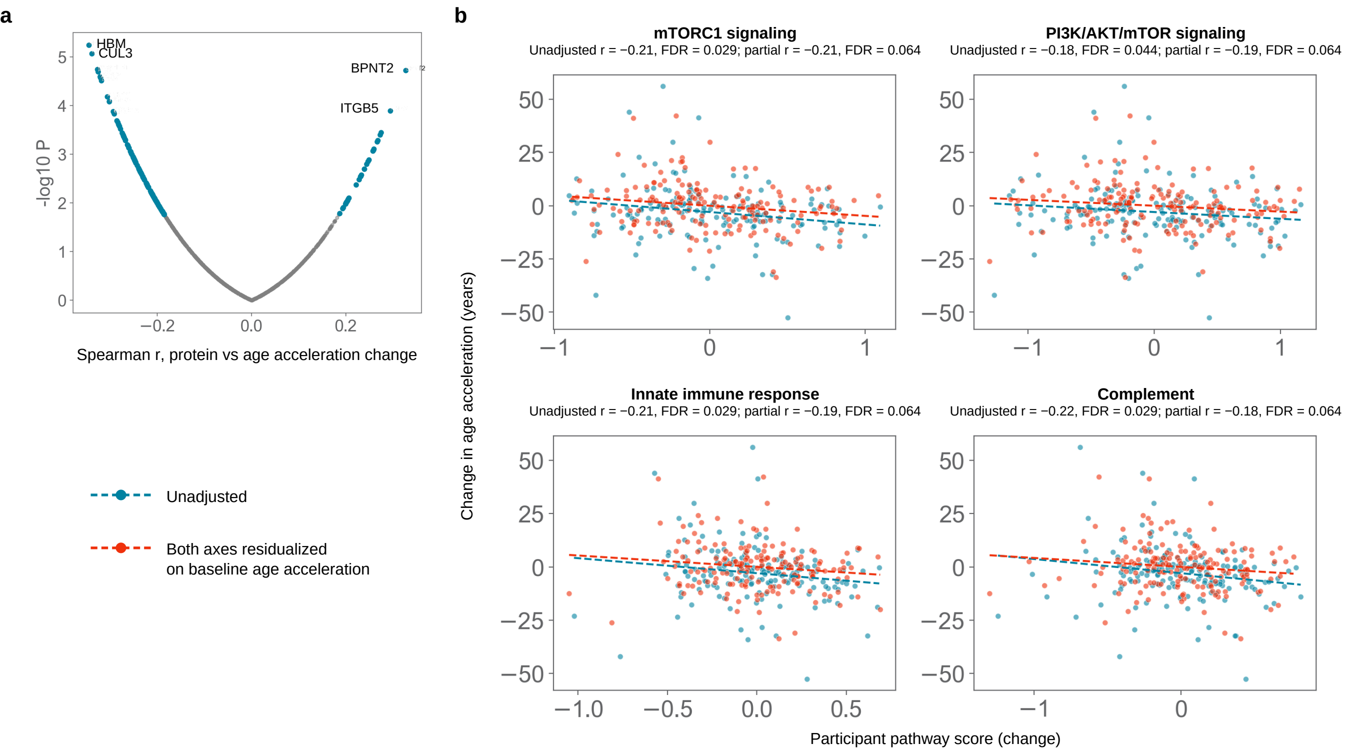
a
HBM
CUL3
BPNT2
ITGB5
-log10 P
Spearman r, protein vs age acceleration change
Unadjusted
Both axes residualized
on baseline age acceleration
b
mTORC1 signaling
Unadjusted r = −0.21, FDR = 0.029; partial r = −0.21, FDR = 0.064
PI3K/AKT/mTOR signaling
Unadjusted r = −0.18, FDR = 0.044; partial r = −0.19, FDR = 0.064
Innate immune response
Unadjusted r = −0.21, FDR = 0.029; partial r = −0.19, FDR = 0.064
Complement
Unadjusted r = −0.22, FDR = 0.029; partial r = −0.18, FDR = 0.064
Change in age acceleration (years)
Participant pathway score (change)

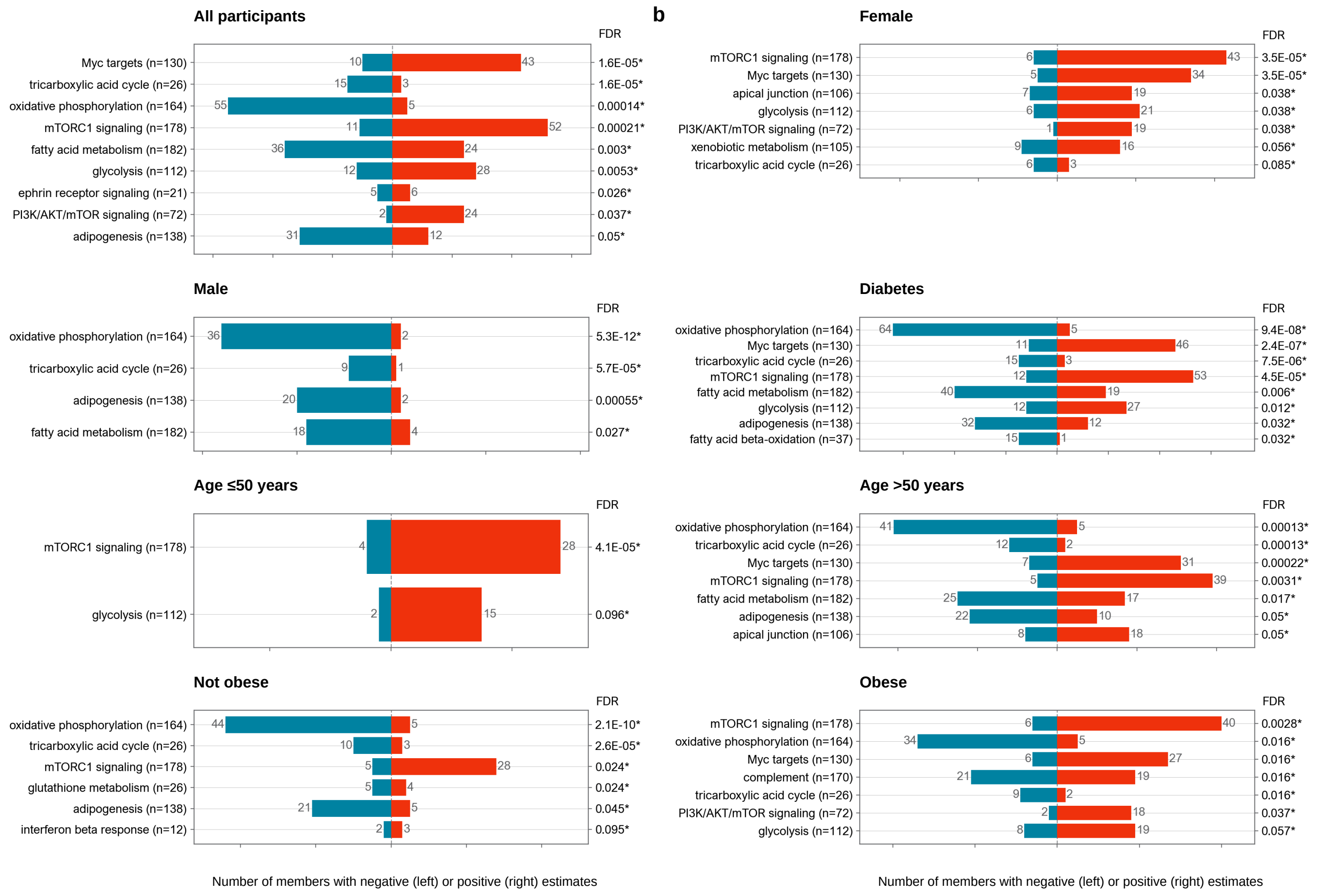
All participants
FDR
Myc targets (n=130) 10 43 1.6E-05*
tricarboxylic acid cycle (n=26) 15 3 1.6E-05*
oxidative phosphorylation (n=164) 55 5 0.00014*
mTORC1 signaling (n=178) 11 52 0.00021*
fatty acid metabolism (n=182) 36 24 0.003*
glycolysis (n=112) 12 28 0.0053*
ephrin receptor signaling (n=21) 5 6 0.026*
PI3K/AKT/mTOR signaling (n=72) 2 24 0.037*
adipogenesis (n=138) 31 12 0.05*
b
Female
FDR
mTORC1 signaling (n=178) 6 43 3.5E-05*
Myc targets (n=130) 5 34 3.5E-05*
apical junction (n=106) 7 19 0.038*
glycolysis (n=112) 6 21 0.038*
PI3K/AKT/mTOR signaling (n=72) 1 19 0.038*
xenobiotic metabolism (n=105) 9 16 0.056*
tricarboxylic acid cycle (n=26) 6 3 0.085*
Male
FDR
oxidative phosphorylation (n=164) 36 2 5.3E-12*
tricarboxylic acid cycle (n=26) 9 1 5.7E-05*
adipogenesis (n=138) 20 2 0.00055*
fatty acid metabolism (n=182) 18 4 0.027*
Diabetes
FDR
oxidative phosphorylation (n=164) 64 5 9.4E-08*
Myc targets (n=130) 11 46 2.4E-07*
tricarboxylic acid cycle (n=26) 15 3 7.5E-06*
mTORC1 signaling (n=178) 12 53 4.5E-05*
fatty acid metabolism (n=182) 40 19 0.006*
glycolysis (n=112) 12 27 0.012*
adipogenesis (n=138) 32 12 0.032*
fatty acid beta-oxidation (n=37) 15 1 0.032*
Age ≤50 years
FDR
mTORC1 signaling (n=178) 4 28 4.1E-05*
glycolysis (n=112) 2 15 0.096*
Age >50 years
FDR
oxidative phosphorylation (n=164) 41 5 0.00013*
tricarboxylic acid cycle (n=26) 12 2 0.00013*
Myc targets (n=130) 7 31 0.00022*
mTORC1 signaling (n=178) 5 39 0.0031*
fatty acid metabolism (n=182) 25 17 0.017*
adipogenesis (n=138) 22 10 0.05*
apical junction (n=106) 8 18 0.05*
Not obese
FDR
oxidative phosphorylation (n=164) 44 5 2.1E-10*
tricarboxylic acid cycle (n=26) 10 3 2.6E-05*
mTORC1 signaling (n=178) 5 28 0.024*
glutathione metabolism (n=26) 5 4 0.024*
adipogenesis (n=138) 21 5 0.045*
interferon beta response (n=12) 2 3 0.095*
Obese
FDR
mTORC1 signaling (n=178) 6 40 0.0028*
oxidative phosphorylation (n=164) 34 5 0.016*
Myc targets (n=130) 6 27 0.016*
complement (n=170) 21 19 0.016*
tricarboxylic acid cycle (n=26) 9 2 0.016*
PI3K/AKT/mTOR signaling (n=72) 2 18 0.037*
glycolysis (n=112) 8 19 0.057*
Number of members with negative (left) or positive (right) estimates
Number of members with negative (left) or positive (right) estimates

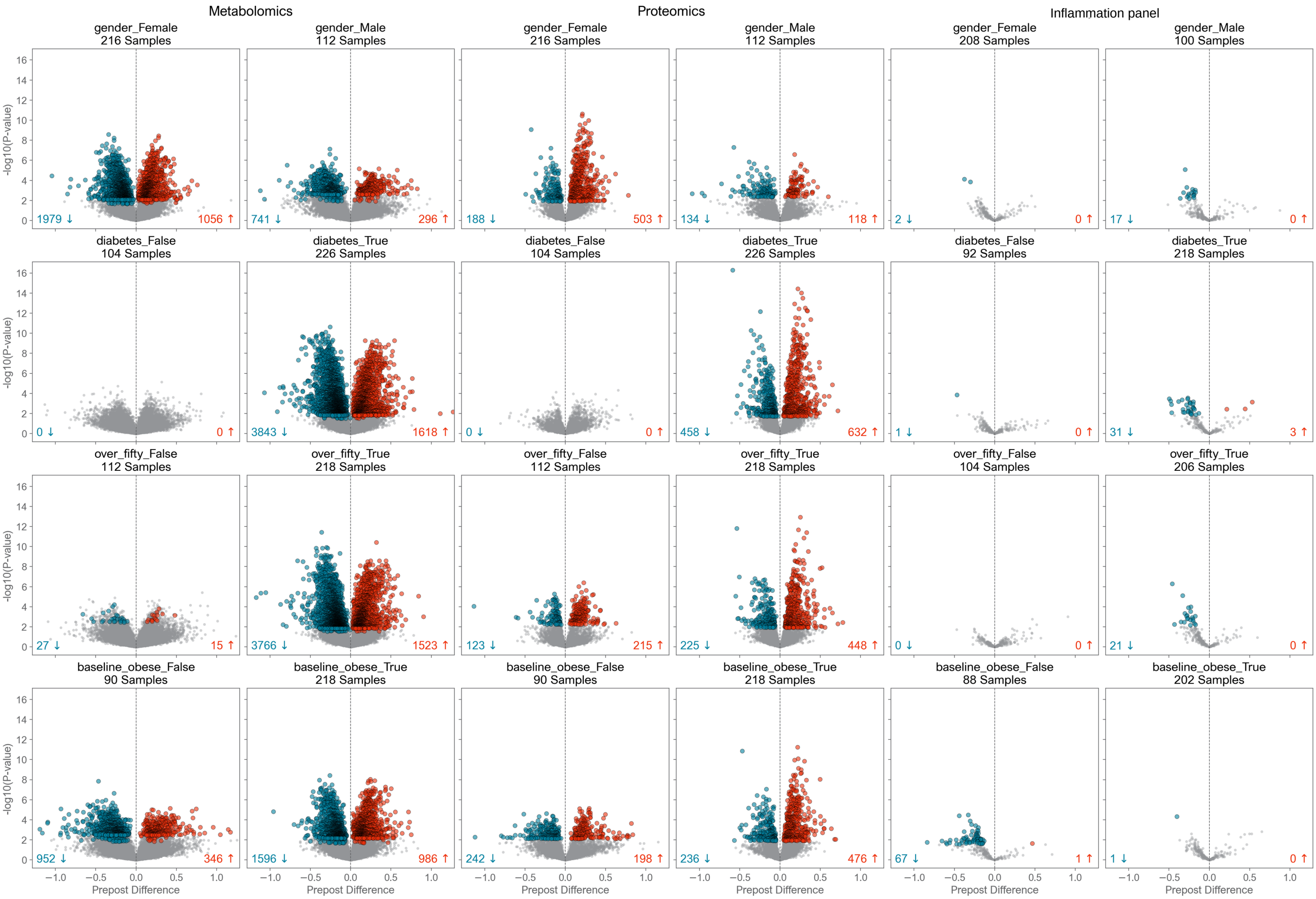

Metabolomics
Proteomics
Inflammation panel
gender_Female
216 Samples
gender_Male
112 Samples
gender_Female
216 Samples
gender_Male
112 Samples
gender_Female
208 Samples
gender_Male
100 Samples
1979 ↓
1056 ↑
741 ↓
296 ↑
188 ↓
503 ↑
134 ↓
118 ↑
2 ↓
0 ↑
17 ↓
0 ↑
diabetes_False
104 Samples
diabetes_True
226 Samples
diabetes_False
104 Samples
diabetes_True
226 Samples
diabetes_False
92 Samples
diabetes_True
218 Samples
0 ↓
0 ↑
3843 ↓
1618 ↑
0 ↓
0 ↑
458 ↓
632 ↑
1 ↓
0 ↑
31 ↓
3 ↑
over_fifty_False
112 Samples
over_fifty_True
218 Samples
over_fifty_False
112 Samples
over_fifty_True
218 Samples
over_fifty_False
104 Samples
over_fifty_True
206 Samples
27 ↓
15 ↑
3766 ↓
1523 ↑
123 ↓
215 ↑
225 ↓
448 ↑
0 ↓
0 ↑
21 ↓
0 ↑
baseline_obese_False
90 Samples
baseline_obese_True
218 Samples
baseline_obese_False
90 Samples
baseline_obese_True
218 Samples
baseline_obese_False
88 Samples
baseline_obese_True
202 Samples
952 ↓
346 ↑
1596 ↓
986 ↑
242 ↓
198 ↑
236 ↓
476 ↑
67 ↓
1 ↑
1 ↓
0 ↑
-log10(P-value)
Prepost Difference

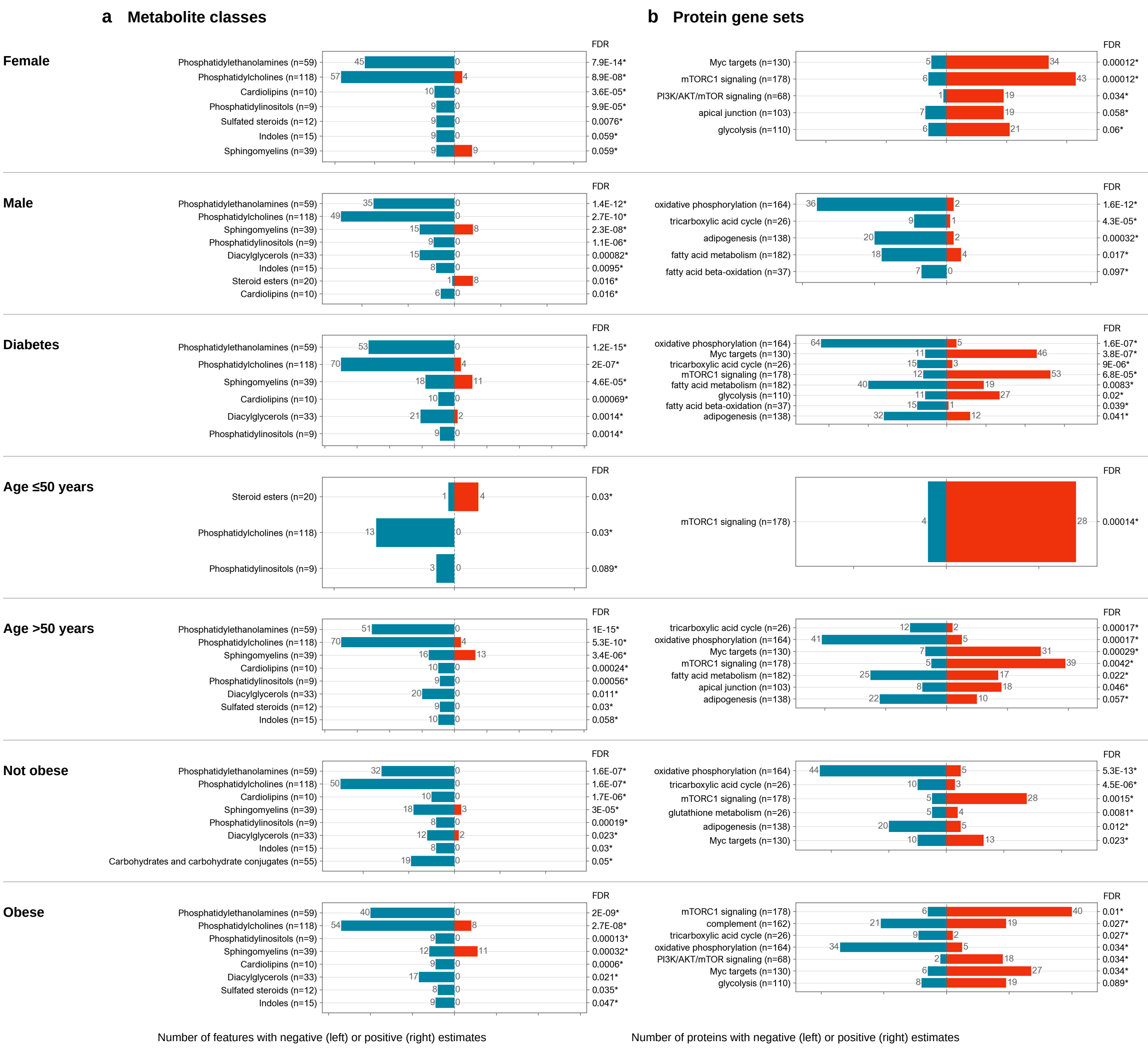

a Metabolite classes
b Protein gene sets
FDR
Female
Phosphatidylethanolamines (n=59) 45 0 7.9E-14*
Phosphatidylcholines (n=118) 57 4 8.9E-08*
Cardiolipins (n=10) 10 0 3.6E-05*
Phosphatidylinositols (n=9) 9 0 9.9E-05*
Sulfated steroids (n=12) 9 0 0.0076*
Indoles (n=15) 9 0 0.059*
Sphingomyelins (n=39) 9 9 0.059*
Myc targets (n=130) 5 34 0.00012*
mTORC1 signaling (n=178) 6 43 0.00012*
PI3K/AKT/mTOR signaling (n=68) 1 19 0.034*
apical junction (n=103) 7 19 0.058*
glycolysis (n=110) 6 21 0.06*
Male
Phosphatidylethanolamines (n=59) 35 0 1.4E-12*
Phosphatidylcholines (n=118) 49 0 2.7E-10*
Sphingomyelins (n=39) 15 8 2.3E-08*
Phosphatidylinositols (n=9) 9 0 1.1E-06*
Diacylglycerols (n=33) 15 0 0.00082*
Indoles (n=15) 8 0 0.0095*
Steroid esters (n=20) 1 8 0.016*
Cardiolipins (n=10) 6 0 0.016*
oxidative phosphorylation (n=164) 36 2 1.6E-12*
tricarboxylic acid cycle (n=26) 9 1 4.3E-05*
adipogenesis (n=138) 20 2 0.00032*
fatty acid metabolism (n=182) 18 4 0.017*
fatty acid beta-oxidation (n=37) 7 0 0.097*
Diabetes
Phosphatidylethanolamines (n=59) 53 0 1.2E-15*
Phosphatidylcholines (n=118) 70 4 2E-07*
Sphingomyelins (n=39) 18 11 4.6E-05*
Cardiolipins (n=10) 10 0 0.00069*
Diacylglycerols (n=33) 21 2 0.0014*
Phosphatidylinositols (n=9) 9 0 0.0014*
oxidative phosphorylation (n=164) 64 5 1.6E-07*
Myc targets (n=130) 11 46 3.8E-07*
tricarboxylic acid cycle (n=26) 15 3 9E-06*
mTORC1 signaling (n=178) 12 53 6.8E-05*
fatty acid metabolism (n=182) 40 19 0.0083*
glycolysis (n=110) 11 27 0.02*
fatty acid beta-oxidation (n=37) 15 1 0.039*
adipogenesis (n=138) 32 12 0.041*
Age ≤50 years
Steroid esters (n=20) 1 4 0.03*
Phosphatidylcholines (n=118) 13 0 0.03*
Phosphatidylinositols (n=9) 3 0 0.089*
mTORC1 signaling (n=178) 4 28 0.00014*
Age >50 years
Phosphatidylethanolamines (n=59) 51 0 1E-15*
Phosphatidylcholines (n=118) 70 4 5.3E-10*
Sphingomyelins (n=39) 16 13 3.4E-06*
Cardiolipins (n=10) 10 0 0.00024*
Phosphatidylinositols (n=9) 9 0 0.00056*
Diacylglycerols (n=33) 20 0 0.011*
Sulfated steroids (n=12) 9 0 0.03*
Indoles (n=15) 10 0 0.058*
tricarboxylic acid cycle (n=26) 12 2 0.00017*
oxidative phosphorylation (n=164) 41 5 0.00017*
Myc targets (n=130) 7 31 0.00029*
mTORC1 signaling (n=178) 5 39 0.0042*
fatty acid metabolism (n=182) 25 17 0.022*
apical junction (n=103) 8 18 0.046*
adipogenesis (n=138) 22 10 0.057*
Not obese
Phosphatidylethanolamines (n=59) 32 0 1.6E-07*
Phosphatidylcholines (n=118) 50 0 1.6E-07*
Cardiolipins (n=10) 10 0 1.7E-06*
Sphingomyelins (n=39) 18 3 3E-05*
Phosphatidylinositols (n=9) 8 0 0.00019*
Diacylglycerols (n=33) 12 2 0.023*
Indoles (n=15) 8 0 0.03*
Carbohydrates and carbohydrate conjugates (n=55) 19 0 0.05*
oxidative phosphorylation (n=164) 44 5 5.3E-13*
tricarboxylic acid cycle (n=26) 10 3 4.5E-06*
mTORC1 signaling (n=178) 5 28 0.0015*
glutathione metabolism (n=26) 5 4 0.0081*
adipogenesis (n=138) 20 5 0.012*
Myc targets (n=130) 10 13 0.023*
Obese
Phosphatidylethanolamines (n=59) 40 0 2E-09*
Phosphatidylcholines (n=118) 54 8 2.7E-08*
Phosphatidylinositols (n=9) 9 0 0.00013*
Sphingomyelins (n=39) 12 11 0.00032*
Cardiolipins (n=10) 9 0 0.0006*
Diacylglycerols (n=33) 17 0 0.021*
Sulfated steroids (n=12) 8 0 0.035*
Indoles (n=15) 9 0 0.047*
mTORC1 signaling (n=178) 6 40 0.01*
complement (n=162) 21 19 0.027*
tricarboxylic acid cycle (n=26) 9 2 0.027*
oxidative phosphorylation (n=164) 34 5 0.034*
PI3K/AKT/mTOR signaling (n=68) 2 18 0.034*
Myc targets (n=130) 6 27 0.034*
glycolysis (n=110) 8 19 0.089*
Number of features with negative (left) or positive (right) estimates
Number of proteins with negative (left) or positive (right) estimates

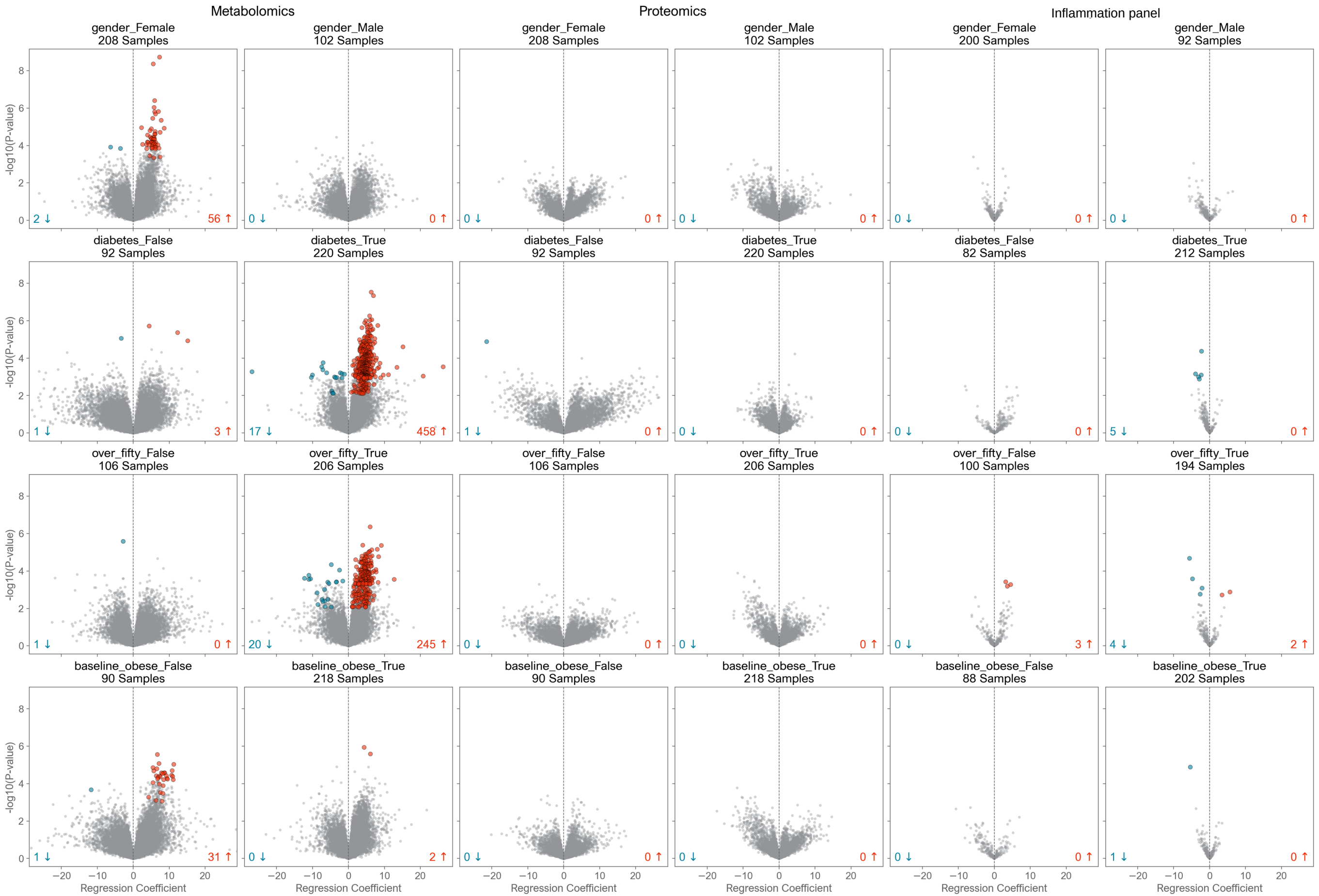

Metabolomics
Proteomics
Inflammation panel
gender_Female 208 Samples
gender_Male 102 Samples
gender_Female 208 Samples
gender_Male 102 Samples
gender_Female 200 Samples
gender_Male 92 Samples
2 ↓
56 ↑
0 ↓
0 ↑
0 ↓
0 ↑
0 ↓
0 ↑
0 ↓
0 ↑
0 ↓
0 ↑
diabetes_False 92 Samples
diabetes_True 220 Samples
diabetes_False 92 Samples
diabetes_True 220 Samples
diabetes_False 82 Samples
diabetes_True 212 Samples
1 ↓
3 ↑
17 ↓
458 ↑
1 ↓
0 ↑
0 ↓
0 ↑
0 ↓
0 ↑
5 ↓
0 ↑
over_fifty_False 106 Samples
over_fifty_True 206 Samples
over_fifty_False 106 Samples
over_fifty_True 206 Samples
over_fifty_False 100 Samples
over_fifty_True 194 Samples
1 ↓
0 ↑
20 ↓
245 ↑
0 ↓
0 ↑
0 ↓
0 ↑
0 ↓
3 ↑
4 ↓
2 ↑
baseline_obese_False 90 Samples
baseline_obese_True 218 Samples
baseline_obese_False 90 Samples
baseline_obese_True 218 Samples
baseline_obese_False 88 Samples
baseline_obese_True 202 Samples
1 ↓
31 ↑
0 ↓
2 ↑
0 ↓
0 ↑
0 ↓
0 ↑
0 ↓
0 ↑
1 ↓
0 ↑
-log10(P-value)
Regression Coefficient

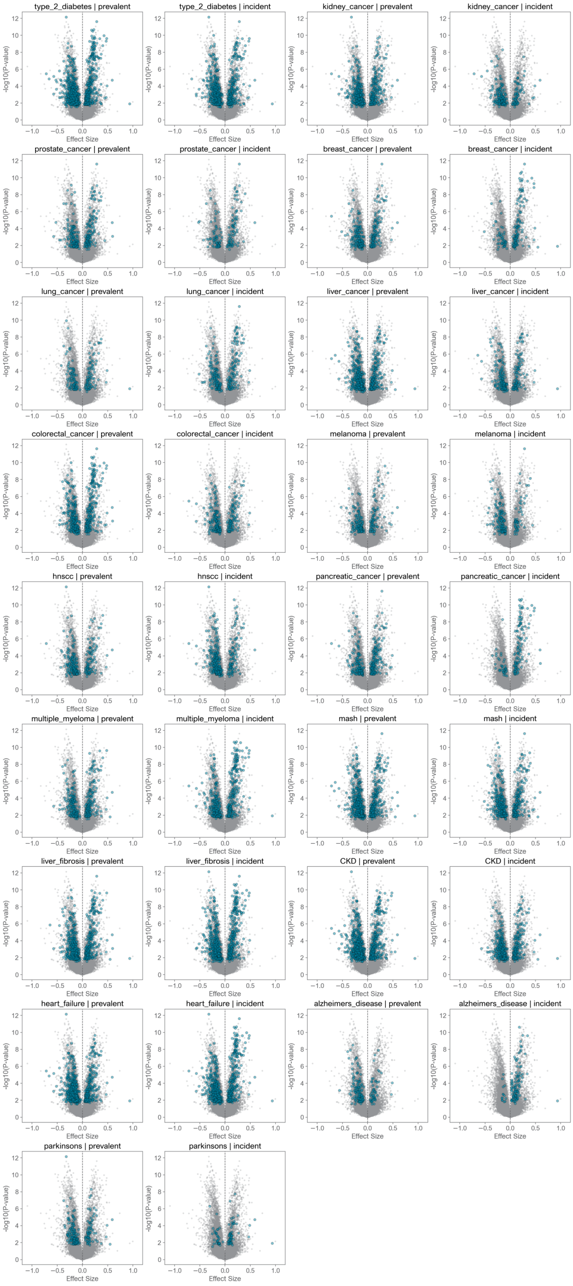
type_2_diabetes | prevalent
type_2_diabetes | incident
kidney_cancer | prevalent
kidney_cancer | incident
prostate_cancer | prevalent
prostate_cancer | incident
breast_cancer | prevalent
breast_cancer | incident
lung_cancer | prevalent
lung_cancer | incident
liver_cancer | prevalent
liver_cancer | incident
colorectal_cancer | prevalent
colorectal_cancer | incident
melanoma | prevalent
melanoma | incident
hnscc | prevalent
hnscc | incident
pancreatic_cancer | prevalent
pancreatic_cancer | incident
multiple_myeloma | prevalent
multiple_myeloma | incident
mash | prevalent
mash | incident
liver_fibrosis | prevalent
liver_fibrosis | incident
CKD | prevalent
CKD | incident
heart_failure | prevalent
heart_failure | incident
alzheimers_disease | prevalent
alzheimers_disease | incident
parkinsons | prevalent
parkinsons | incident
-log10(P-value)
0
2
4
6
8
10
12
Effect Size
−1.0
−0.5
0.0
0.5
1.0

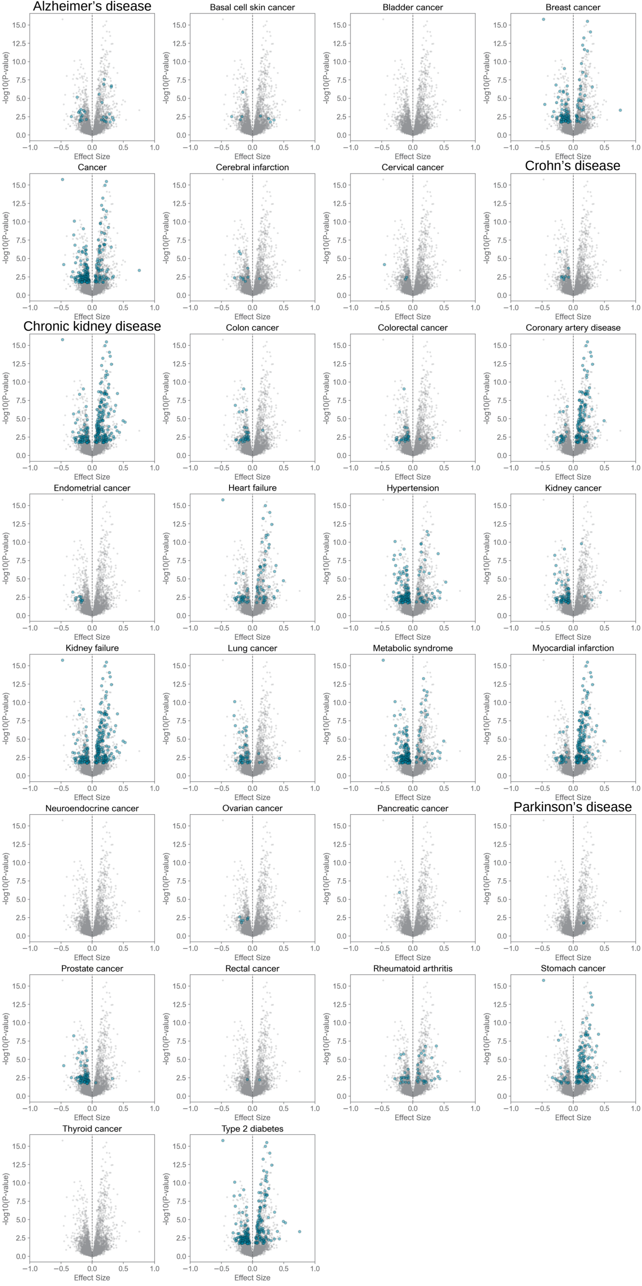
Alzheimer’s disease
Basal cell skin cancer
Bladder cancer
Breast cancer
Cancer
Cerebral infarction
Cervical cancer
Crohn’s disease
Chronic kidney disease
Colon cancer
Colorectal cancer
Coronary artery disease
Endometrial cancer
Heart failure
Hypertension
Kidney cancer
Kidney failure
Lung cancer
Metabolic syndrome
Myocardial infarction
Neuroendocrine cancer
Ovarian cancer
Pancreatic cancer
Parkinson’s disease
Prostate cancer
Rectal cancer
Rheumatoid arthritis
Stomach cancer
Thyroid cancer
Type 2 diabetes
-log10(P-value)
Effect Size

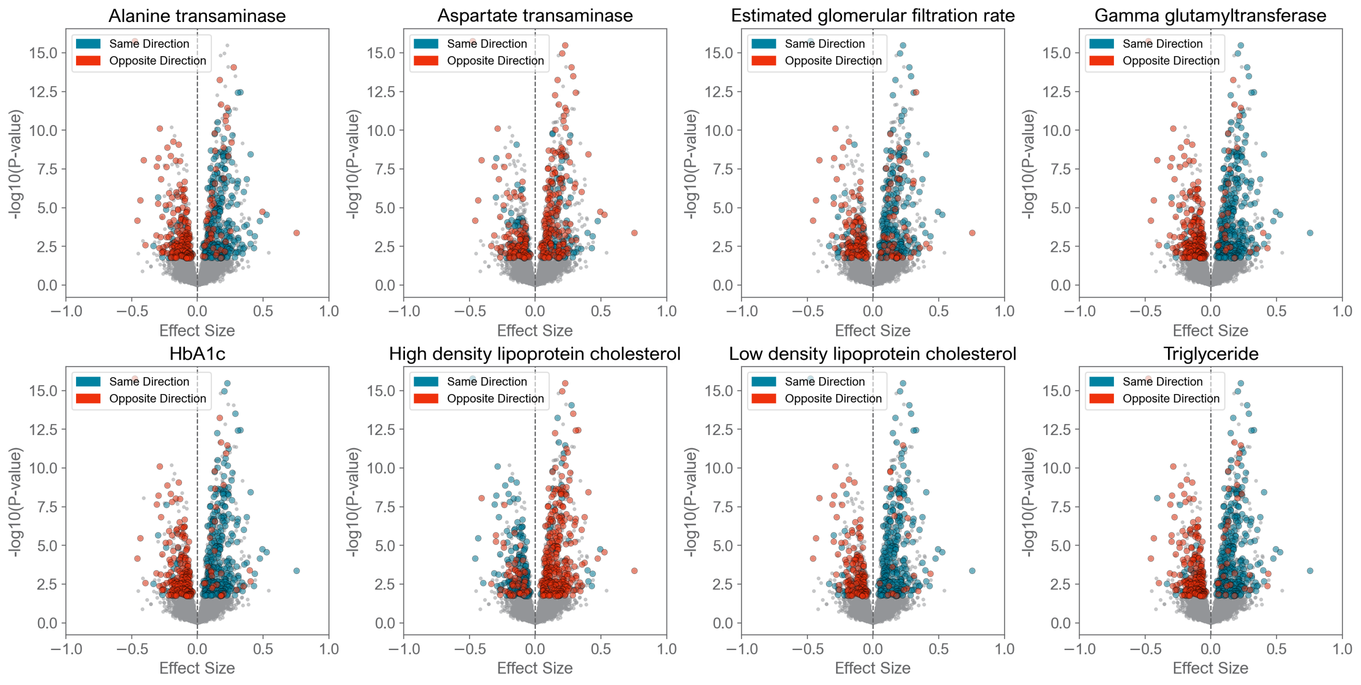

Alanine transaminase
Aspartate transaminase
Estimated glomerular filtration rate
Gamma glutamyltransferase
HbA1c
High density lipoprotein cholesterol
Low density lipoprotein cholesterol
Triglyceride
Same Direction
Opposite Direction
-log10(P-value)
Effect Size
0.0
2.5
5.0
7.5
10.0
12.5
15.0
−1.0
−0.5
0.0
0.5
1.0

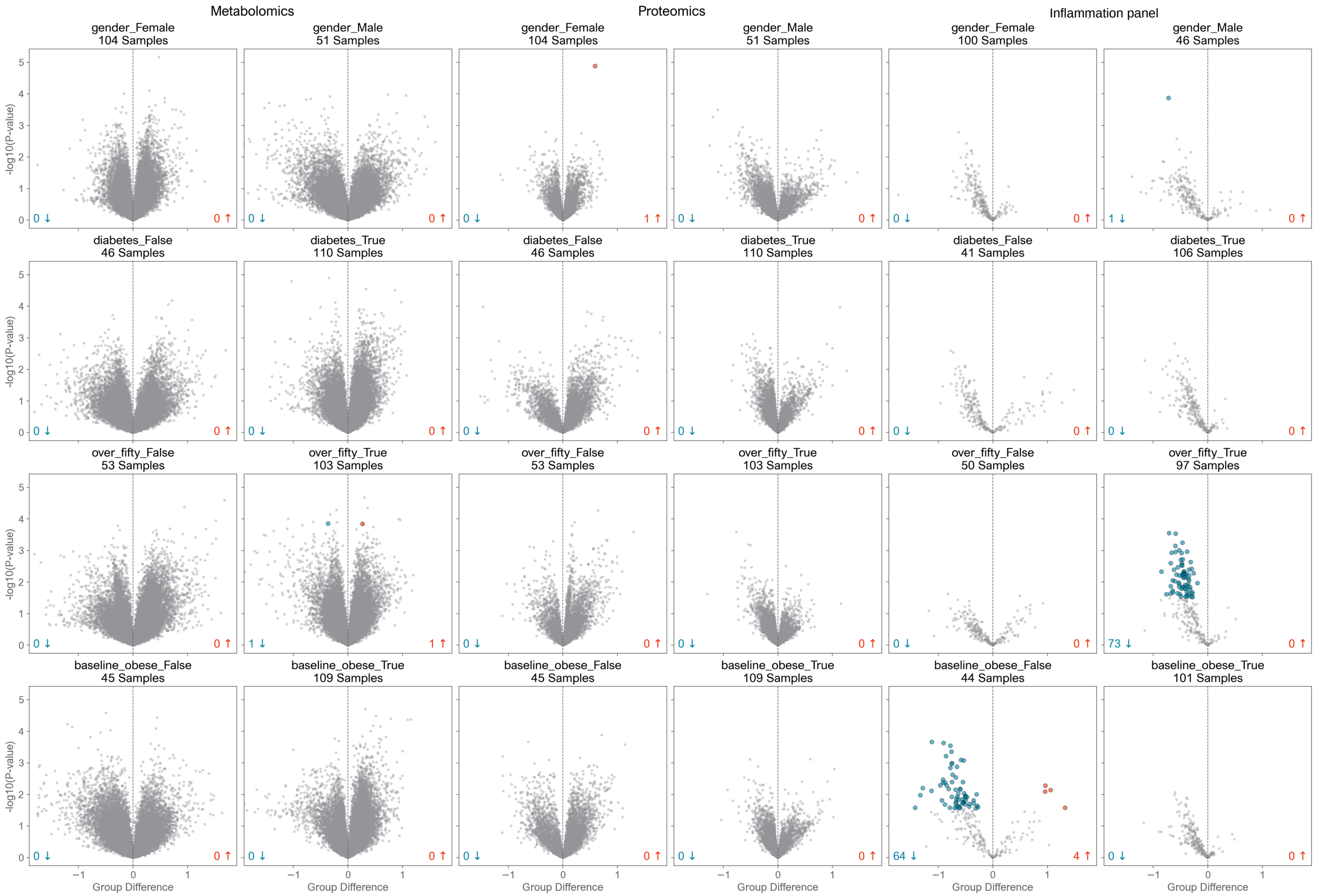
Metabolomics
Proteomics
Inflammation panel
gender_Female
104 Samples
gender_Male
51 Samples
gender_Female
104 Samples
gender_Male
51 Samples
gender_Female
100 Samples
gender_Male
46 Samples
0 ↓ 0 ↑
0 ↓ 0 ↑
0 ↓ 1 ↑
0 ↓ 0 ↑
0 ↓ 0 ↑
1 ↓ 0 ↑
diabetes_False
46 Samples
diabetes_True
110 Samples
diabetes_False
46 Samples
diabetes_True
110 Samples
diabetes_False
41 Samples
diabetes_True
106 Samples
0 ↓ 0 ↑
0 ↓ 0 ↑
0 ↓ 0 ↑
0 ↓ 0 ↑
0 ↓ 0 ↑
0 ↓ 0 ↑
over_fifty_False
53 Samples
over_fifty_True
103 Samples
over_fifty_False
53 Samples
over_fifty_True
103 Samples
over_fifty_False
50 Samples
over_fifty_True
97 Samples
0 ↓ 0 ↑
1 ↓ 1 ↑
0 ↓ 0 ↑
0 ↓ 0 ↑
0 ↓ 0 ↑
73 ↓ 0 ↑
baseline_obese_False
45 Samples
baseline_obese_True
109 Samples
baseline_obese_False
45 Samples
baseline_obese_True
109 Samples
baseline_obese_False
44 Samples
baseline_obese_True
101 Samples
0 ↓ 0 ↑
0 ↓ 0 ↑
0 ↓ 0 ↑
0 ↓ 0 ↑
64 ↓ 4 ↑
0 ↓ 0 ↑
-log10(P-value)
Group Difference
0
1
2
3
4
5
−1
0
1

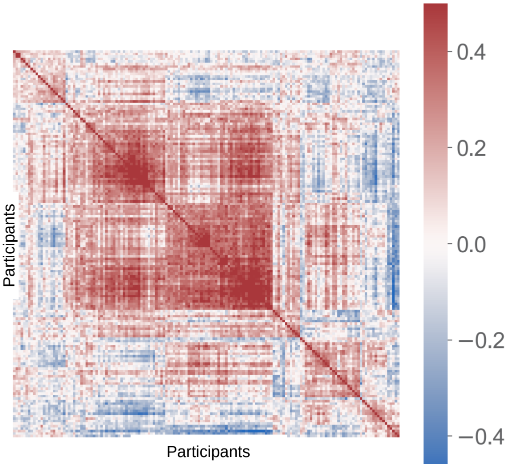
Participants
Participants
0.4
0.2
0.0
−0.2
−0.4
Spearman r
