## Supplementary Methods for "Real-world longitudinal multi-omics separates weight-loss-associated responses to incretin agonists from a broader response not dependent on weight loss"

Address for correspondence:

**Supplementary Methods**

**Study cohort and clinical data**

***Cohort definition and sample selection***

The study cohort was defined by querying the Sapient DynamiQ biorepository for participants prescribed an incretin agonist (semaglutide, liraglutide, dulaglutide or tirzepatide) for at least 90 days. The cohort was further restricted to participants with plasma samples collected both before and during treatment. For each participant, we selected the sample collected closest to, but before, the first recorded prescription date (pre-treatment sample) and the sample collected closest to, but before, the last recorded prescription date (post-treatment sample), together with the associated clinical metadata.

***Weight change and response groups***

Weight measurements were taken from linked electronic health records (EHR). For the pre-treatment time point, we used the weight measurement closest to, but before, the initial prescription date. For the post-treatment time point, we used the weight measurement closest to, but before, the last known prescription date, provided it was recorded within 30 days of sample collection. Percent weight change was calculated between these two measurements. Participants were classified as having no weight loss (weight change ≥ 0%), weight loss of 0--5%, or weight loss of 5% or more.

***Baseline obesity***

Baseline obesity was defined using the body mass index (BMI) recorded closest to, but before, treatment initiation, with BMI ≥ 30 kg m⁻² classified as obese.

***Clinical laboratory measures***

High-density lipoprotein cholesterol, low-density lipoprotein cholesterol, triglycerides, glycated hemoglobin (HbA1c) and estimated glomerular filtration rate (eGFR) were taken from EHR records. The pre-treatment value was the measurement closest to treatment initiation, provided it was recorded between 90 days before and 30 days after initiation. The post-treatment value was the measurement closest to the end of treatment, provided it was recorded between 30 days before and 90 days after the end of treatment.

**Mass spectrometry-based plasma proteomics**

***Materials***

Lyophilized HeLa digest standard (Pierce) was used to assess system suitability before injection of study samples. Acetonitrile, water, ammonium bicarbonate, tris(2-carboxyethyl)phosphine (TCEP), iodoacetamide (IAA), formic acid, sodium deoxycholate (SDC) and Trizma base were purchased from Fisher Scientific (Waltham, MA, USA) and were LC-MS grade where applicable. Lys-C and trypsin were purchased from Thermo Fisher Scientific (Waltham, MA, USA). Evosep C18 Pure Evotips and the Evosep 8 cm × 150 µm column were purchased from Evosep Biosystems (Odense, Denmark).

***Sample handling***

K2-EDTA plasma samples were stored at −80 °C and transported on dry ice to the central laboratory (Sapient Bioanalytics, San Diego, CA, USA). Samples were thawed over 10 min using a custom rapid plate-thaw system and placed on an orbital shaker at 550 rpm and 4 °C for 10 min to ensure a uniform sample matrix.

***Sample preparation***

Plasma (12.5 µL) was combined with 1% LMNG and binding buffer (5× v/v) and incubated with Sapient's protein-enrichment magnetic nanoparticles for 30 min at room temperature with gentle vortex mixing. Nanoparticles were washed three times with water to remove non-specifically bound proteins. Disulfide bonds were reduced with 5 mM TCEP in 50 mM ammonium bicarbonate, pH 8.0, for 45 min at 45 °C with gentle mixing, and samples were then cooled to room temperature. Free cysteines were alkylated with 15 mM IAA for 30 min at room temperature in the dark, and the reaction was quenched with 15 mM dithiothreitol for 15 min at room temperature. Nanoparticles were washed with ammonium bicarbonate, pH 8.0, to remove excess reagents, and proteins were digested on-bead overnight at 37 °C in ammonium bicarbonate, pH 8.0, containing 1% SDC, 300 ng trypsin and 100 ng Lys-C per sample. After digestion, samples were acidified with 3% formic acid and filtered through a 0.2 µm PVDF filter plate.

***Data acquisition***

Tryptic peptides (800 ng) were loaded onto Evosep C18 Pure Evotips, separated using the Evosep One 60 samples-per-day (24-min) preformed gradient, and analyzed on a Bruker timsTOF HT mass spectrometer operating in DIA-PASEF mode. Settings were: positive ion mode; m/z 100--1,700 scan range; 1/K₀ ramp of 0.70--1.30 with a 60 ms ramp time; and collision energy ramped from 20 eV at 1/K₀ 0.60 to 65 eV at 1/K₀ 1.60. DIA was performed with a 12 × 2 asymmetric window scheme covering m/z 300--1,350 and 1/K₀ 0.70--1.30.

***Data processing and quality control***

Data were analyzed in Spectronaut 20 using the directDIA workflow against a project-specific gas-phase fractionation spectral library and the UniProt human canonical plus common isoforms FASTA (42,490 entries). Protein and peptide false discovery rates were set at 1%, with peptide lengths of 6--52 residues and up to two missed cleavages. Carbamidomethylation of cysteine was the only fixed modification; variable modifications were protein N-terminal acetylation, deamidation (N, Q), methionine oxidation and conversion of N-terminal glutamine to pyroglutamate. Data were normalized using MaxLFQ^1^ before export. System suitability was established before sample injection by triplicate injection of the HeLa digest standard. Instrument stability and preparation reproducibility were monitored using bracketing samples of pooled plasma, prepared in independent wells and injected at the beginning and end of each run and every 12 h in between.

**Untargeted metabolomics**

***Sample handling***

Samples were transported on dry ice to the central laboratory, randomized on dry ice on receipt and stored at −80 °C. Samples were de-identified before receipt, and all analyses were performed blinded.

***Metabolite extraction***

Samples were thawed over 4 min using a custom rapid plate-thaw system and placed on an orbital shaker at 550 rpm and 4 °C for 10 min. In a shallow 96-well plate, plasma was transferred into acidified acetonitrile and methanol containing internal standards (¹³C-, ¹⁵N- and deuterium-labeled polar molecules and lipids spanning the full retention-time range, from Cayman Chemical, Sigma-Aldrich and Avanti Polar Lipids). Samples were shaken at 550 rpm and 4 °C for 10 min and centrifuged at 6,000 g and 4 °C for 10 min. For positive ion mode, 2 µL of supernatant was added to 148 µL of water:methanol (80:20) containing 0.05% formic acid and 1 mM ammonium formate. For negative ion mode, 15 µL of supernatant was added to 135 µL of water:methanol (60:40) containing 0.1% acetic acid. Both dilution solutions contained additional deuterium-labeled internal standards.

***Rapid LC-MS analysis***

Samples were analyzed by rapid liquid chromatography-mass spectrometry (rLC-MS)^2^, which uses rapid nano-valve switching with a three-pump system delivering a multi-tiered isocratic elution, coupled to a Bruker timsTOF Pro 2 mass spectrometer (Bruker, Billerica, MA, USA). Samples were injected onto a custom-packed silica-based mixed-mode column supporting both reversed-phase and weak ion-pairing retention. Data were acquired in positive and negative electrospray ionization modes with an injection-to-injection cycle time of 52 s. Elution began with methanol:water (20:80) containing 0.05% acetic acid and 2.5 mM ammonium acetate (positive mode) or methanol:water (35:65) containing 0.02% acetic acid and 5 mM ammonium acetate (negative mode), and ended with methanol:isopropanol (80:20) containing 0.1% acetic acid and 7.5 mM ammonium acetate (positive mode) or methanol:isopropanol (70:30) containing 0.1% acetic acid and 20 mM ammonium acetate (negative mode). Mass spectrometer settings were: dry gas temperature 350 °C; dry gas flow 8 L min⁻¹ (positive) and 6 L min⁻¹ (negative); nebulizer 50 psi; sheath gas temperature 150 °C (positive) and 250 °C (negative); sheath gas flow 3 L min⁻¹; capillary voltage 3,500 V (positive) and 3,000 V (negative); end plate offset 2,000 V (positive) and 1,500 V (negative); mass range m/z 50--1,700; and acquisition at 9 spectra s⁻¹.

***Run quality control***

For each 384-well plate, the isotopically labeled internal standards added at each preparation step were monitored for stability, retention time and mass accuracy. Pre-aliquoted commercial pooled plasma (BioIVT, Westbury, NY, USA) was placed in wells A1, D12 and H12 of each 96-well sample plate and prepared identically to study samples as an internal bracketing control. A separate aliquot of pooled EDTA plasma was prepared by hand and injected as an external control, together with system blanks to assess background.

***Data extraction and normalization***

After chromatographic drift correction using landmark features spanning the dataset^2^, metabolite features were extracted using custom deep-learning-based feature detection. Features were then normalized for plate-to-plate variation by batch median normalization.

***Metabolite annotation***

Extracted features were matched against Sapient's internal library of commercial reference standards. Protonated and deprotonated ions and observed adducts were retained, and isotopes, adducts and fragments were flagged alongside their parent ions. Adducts were identified by comparing observed m/z differences with a curated list of common adducts (for example, [M+H]⁺ and [M+Na]⁺), applying polarity-specific filters where relevant and requiring a retention-time difference of ≤ 0.05 min and a mass accuracy of ±10 ppm. Candidate parent--adduct relationships were retained only when the sample-wise Pearson correlation of peak intensities exceeded 0.70 (P ≤ 0.05). Isotopologues were identified by estimating the number of carbon atoms in each parent ion from its m/z using a model derived from the Human Metabolome Database^3^, calculating the expected ¹³C abundance ratio (1.1% per carbon), and confirming candidate peaks whose observed intensity ratios and retention-time proximity (≤ 0.05 min) matched these predictions. Fragments (for example, losses of H₂O or CO₂) were annotated by comparing observed m/z differences with predefined fragmentation rules. In the final dataset, each parent ion was grouped with its associated adducts, isotopologues and fragments.

**Inflammation panel (NULISAseq)**

***Sample analysis***

Samples were thawed over 4 min using a custom rapid plate-thaw system and centrifuged at 2,000 g for 10 min, and 25 µL of supernatant was transferred to a 96-well plate for analysis on the ARGO HT system (Alamar Biosciences). The ARGO HT is a self-contained instrument preloaded with all required reagents and buffers, including a deactivation buffer (isopropanol:water, 70:30), a water rinse buffer and a proprietary wash buffer.

Each target is measured with a pair of antibodies pooled into a multiplex panel: a capture antibody conjugated to partially double-stranded DNA containing a poly-A tail and a target-specific molecular identifier, and a detection antibody conjugated to partially double-stranded DNA containing a biotin group and a matching target-specific identifier. Binding of both antibodies to the target antigen forms an immunocomplex, which is captured on paramagnetic oligo-dT beads by dT--polyA hybridization while the sample matrix and unbound detection antibodies are washed away. Because dT--polyA binding is salt-sensitive, the immunocomplexes are then released into a low-salt buffer and recaptured on streptavidin-coated paramagnetic beads, and further washes remove unbound capture antibodies. A ligation mix containing T4 DNA ligase, a DNA ligator sequence and a sample-specific molecular identifier then joins the proximal DNA ends of each antibody pair, generating a reporter molecule that carries the target and sample identifiers and is quantified by next-generation sequencing.

***Sequencing***

The pooled reporter library from each run was sequenced on an Element Biosciences AVITI system with 150-base single-end reads on a dedicated flow cell, at a depth of 500 million reads per library, using custom libraries provided by Alamar Biosciences.

***Quality control***

An exogenous reporter added to each well served as an internal control for normalization. Pooled plasma was included as a sample control to assess run suitability and intra- and inter-plate coefficients of variation. An independent plasma pool served as an inter-plate control, used to normalize samples across runs and to assess assay precision. Assay buffer served as a negative control to determine limits of detection and overall assay performance.

***Data processing***

Sequencing data (FASTQ files) were processed and reviewed in the Alamar Control Center. NULISA protein quantification (NPQ) values were calculated by normalizing each target's reads to the internal control and dividing by the target-specific median of the inter-plate control replicates. Values were multiplied by 10,000, incremented by 1 and log₂-transformed.

**Data preprocessing**

All analyses were performed with an internal analysis package built on Python and scikit-learn^4^, and each data type was preprocessed independently. For metabolomics, isotopologues flagged during feature extraction were confirmed by Spearman correlation between the intensity of the parent peak and the putative isotopologue, and those with a correlation above 0.9 were removed. Features below the limit of detection in more than 20% of samples, and features with intensity below 1,000 arbitrary units, were also removed. For proteomics and the inflammation panel, features below the limit of detection in more than 20% of samples were removed. For all three layers, missing values were imputed feature by feature with values drawn from a uniform distribution between half the feature minimum and the feature minimum, and the data were then log-transformed and standardized. Principal component analysis was used to visualize batch effects and outliers, and preprocessing was repeated with any outlying samples removed.

**Statistical analyses**

***Pharmacodynamic biomarkers***

Features that changed with treatment were identified by two-sided paired t-tests between each participant's pre- and post-treatment samples, in the full cohort and within strata defined by sex, age (above or below 50 years), diabetes status and baseline obesity.

***Association with weight loss***

The association between each feature's log₂ fold change and percent weight change was assessed by Spearman correlation and by ordinary least squares regression adjusted for sex, age, diabetes status and baseline body mass index, in the full cohort and within the same strata.

***Baseline associations with subsequent weight loss***

Two complementary analyses related baseline feature levels to subsequent weight loss. To treat weight loss as continuous, percent weight change was regressed on baseline feature level with age and sex as covariates. To capture features that separate responders, baseline levels were compared between participants who lost more than 5% of body weight and those who did not by two-sided unpaired t-test. Both analyses were repeated within strata.

***Multiple-testing correction***

P values were adjusted by the Benjamini--Hochberg procedure at a false discovery rate (FDR) of 0.1 for each analysis. For metabolomics, annotated and unannotated features were corrected separately.

***Enrichment analysis***

Enrichment was tested separately for metabolite compound classes and for protein and inflammation-panel gene sets. For each class or gene set with at least five measured members, a 2 × 2 contingency table of membership versus significance was tested by one-sided Fisher's exact test, and P values were corrected across all classes or gene sets.

***Disease associations of treatment-responsive features***

For metabolites, we used the DynamiQ biorepository to identify features associated with disease. For each outcome, participants with a diagnosis, defined by ICD-10 code, were identified. Samples collected more than 90 days before the first recorded diagnosis were classified as incident, and samples collected between 90 days before and 5 years after the first diagnosis as prevalent. Cases were matched to healthy controls by the proportion of each sex and age-group combination, and the numbers of cases and controls for each outcome are given in Supplementary Table 16a. Healthy controls were participants never diagnosed with any of the following: cancer (C00--C97), cardiovascular disease (I00--I99), chronic respiratory disease (J40--J47, J60--J70), diabetes (E08--E13), endocrine disorders (E00--E07, E20--E27, E31--E35), metabolic disorders (E70--E72, E74--E79, E80, E83--E85, E88.4), chronic liver disease (B18, K70--K76), renal disease (N17--N19), autoimmune disease (K50--K52, M05, M06.0--M06.39, M32, M34, G35), neurodegenerative disease (G10--G32), neurological disorders (G37, G40, G80--G83), cerebrovascular disease (F01, G46), substance use disorders (F10--F19), muscular dystrophy (G71), tuberculosis (A15--A19), HIV (B20) or post-COVID-19 condition (U09). For each outcome, incident and prevalent analyses were performed separately by logistic regression of case status on each metabolite, adjusting for age and sex where sample size allowed, with FDR correction applied separately to annotated and unannotated features. Fig. 5a reports the incident analysis.

For proteins, we used published UK Biobank proteomic case--control summary statistics^5^, identifying overlaps between proteins associated with each outcome and treatment-responsive proteins, and selecting outcomes equivalent or closely related to those analyzed in DynamiQ.

***Metabolic aging clock***

The metabolic aging clock was retrained following the approach of our previous study^2^ to minimize batch effects. Features selected in the original model were aligned to the present dataset, and healthy controls (defined as in the disease analyses) were drawn from the same analytical batch as the treated cohort and split 70/30 at the participant level into training and held-out validation sets. A scaler fitted to the training set was applied to the validation set and the treated cohort. Lasso hyperparameters were tuned on the training set with Optuna^6^ using five-fold cross-validation. The loss for each fold was the mean absolute error between predicted and chronological age plus a penalty equal to 0.5 times the difference between the test-fold and training-fold mean absolute errors, and Optuna minimized the mean loss across folds. Because healthy controls are on average younger than individuals with disease, predictions were corrected by fitting a linear regression of predicted on chronological age in the training set and applying its inverse (Methods). The model was then evaluated on the held-out validation set before being applied to the treated cohort.

***Pathway response scores and the change in age acceleration***

For each of 12 protein gene sets, a per-participant response score was computed as the mean z-scored log₂ fold change of the gene set's treatment-responsive members. Scores were related to the within-person change in age acceleration by Spearman correlation, by partial Spearman correlation adjusting for baseline age acceleration, and by linear regression including baseline age acceleration as a covariate, with Benjamini--Hochberg correction across gene sets (Supplementary Table 14). Inflammation-panel analytes were analyzed in the same way (Fig. 4h).

***Sample hemolysis index***

A per-sample hemolysis index was calculated as the mean standardized abundance of nine erythrocyte-enriched proteins (HBA1, HBB, HBD, CA1, CA2, PRDX2, CAT, BLVRB and SLC4A1). Its relationship with the within-person change in age acceleration was assessed by correlation (Supplementary Table 11; Supplementary Note 1).

***Correlation network***

Weight-loss-dependent features were defined as treatment-responsive features that were also significantly associated with weight loss. Features not dependent on weight loss were treatment-responsive features whose association with weight loss had P > 0.5 and an absolute estimate below 0.2. Spearman correlation coefficients were calculated between every pair of features, and the network was drawn with edges for |r| > 0.7, showing only nodes with at least one such edge.

**Supplementary Note 1**

***Erythrocyte contamination, sampling interval and the change in metabolic age acceleration***

Plasma proteomic and metabolomic profiles are sensitive to the lysis of erythrocytes during collection and processing, which releases hemoglobin and other intracellular proteins and metabolites into the sample. Because the paired samples in this study were collected in routine care rather than under a common protocol, differences in handling between the pre- and post-treatment draws could alter the measured profiles independently of treatment. We therefore asked whether erythrocyte contamination differed between paired samples and whether it was related to the decrease in metabolic age acceleration reported in the main text.

We estimated erythrocyte contamination with a per-sample hemolysis index, defined as the mean standardized abundance of nine erythrocyte-enriched proteins (HBA1, HBB, HBD, CA1, CA2, PRDX2, CAT, BLVRB and SLC4A1; Supplementary Methods). Four of the nine index proteins were among the proteins that increased between paired samples, namely hemoglobin subunit β (HBB; +0.40 log2 fold change, FDR = 5.4 × 10⁻³), hemoglobin subunit δ (HBD; +0.36, FDR = 0.030), carbonic anhydrase 2 (CA2; +0.27, FDR = 6.9 × 10⁻⁵) and flavin reductase (BLVRB; +0.21, FDR = 0.028) (Supplementary Table 3), indicating modestly higher erythrocyte contamination in post-treatment samples. Storage time before analysis, by contrast, was unrelated to the change in age acceleration for both the pre-treatment sample (r = 0.06, P = 0.46) and the post-treatment sample (r = 0.10, P = 0.18) (Supplementary Table 11).

Erythrocyte contamination was associated with the change in age acceleration. Contamination in the post-treatment sample correlated with the within-person change in age acceleration (r = −0.24, P = 1.9 × 10⁻³), as did the within-person change in contamination (r = −0.23, P = 3.7 × 10⁻³), whereas contamination in the pre-treatment sample did not (r = 0.05, P = 0.54) (Supplementary Table 11). The same four erythrocyte proteins were among the proteins whose change correlated with the change in age acceleration (HBB, Spearman r = −0.25, FDR = 0.023; HBD, r = −0.25, FDR = 0.025; CA2, r = −0.21, FDR = 0.056; BLVRB, r = −0.21, FDR = 0.060; Supplementary Table 13). Participants whose post-treatment sample carried more erythrocyte material therefore tended to show larger decreases in age acceleration, so differential contamination between paired samples could contribute to the decrease reported in the main text.

The decrease in age acceleration was also unrelated to the timing of sample collection. It did not correlate with the interval between paired samples (median 406 days, range 112 to 1,131; Spearman r = −0.05, P = 0.51), with the duration of treatment at the post-treatment sample (median 359 days, range 95 to 1,109; r = −0.06, P = 0.48) or with the calendar date of the pre-treatment sample (r = −0.06, P = 0.46) (Supplementary Table 10). The decrease therefore does not reflect elapsed time between samples or drift in measurement across the collection period.

Together, these analyses indicate that erythrocyte contamination was modestly higher in post-treatment samples and was associated with the change in age acceleration, whereas the change was unrelated to sampling interval, treatment duration, calendar date or storage time. A model adjusting the change in age acceleration for contamination was not part of the present analysis, so a contribution of differential erythrocyte contamination to the decrease cannot be excluded; repeated sampling under a standardized collection protocol would resolve it.
